# Mechanistic basis for targeting homologous recombination defective liver cancer via synthetic lethality

**DOI:** 10.1101/2022.07.21.500952

**Authors:** Ming Zeng, Zizhi Tang, Laifeng Ren, Haibin Wang, Xiaojun Wang, Wenyuan Zhu, Xiaobing Mao, Zeyang Li, Xianming Mo, Jun Chen, Junhong Han, Daochun Kong, Jianguo Ji, Antony M. Carr, Cong Liu

## Abstract

Many cancers harbour homologous recombination defects (HRD). The identification of PARP inhibitors as synthetic lethal with HRD has led to new therapeutic strategies for HRD cancers. Here we report a subtype of HRD that is caused by the perturbation of a previously uncharacterised proteasome variant, CDW19S, in hepatitis virus B (HBV) positive hepatocellular carcinoma (HBVHCC). CDW19S contains the 19S complex decorated with a Cullin 4 ubiquitin ligase (CRL4^WDR70^) that is assembled at broken chromatin and regulates end processing nucleases. The HBV oncoprotein, HBx, prevents integration of the CRL4 backbone into CDW19S. We show that CDW19S directly ubiquitinates ADRM1^Rpn13^, targeting it for degradation, and that HBx interferes with this, leading to the imposition of a novel ADRM1^Rpn13^-dependent resection barrier that results in HRD and promotes carcinogenesis with concurrent *TP53* loss. Using cellular and patient-derived xenograft models we demonstrate that HRD in HBVHCC can be exploited to restrict tumour progression. Our work clarifies the mechanism of a virally-induced HRD and suggests a new route for targeted HBVHCC therapy.

## Introduction

Long-term liver infection with Hepatitis virus B (HBV) predisposes carriers to hepatocellular carcinoma (HCC)^1^. Therapy for late-stage HBVHCC remains a major challenge, with poor therapy responses and low overall survival^2,3^. HBV targets DNA repair via HBx, the viral oncoprotein that antagonizes homologous recombination. HBx subverts the Cullin4-DDB1-RING (CRL4) ubiquitin ligase for viral purposes^4^. Among the CRL4 subcomplexes that subsequently show reduced function is CRL4^WDR70^, which functions in HR and chromatin remodelling^5,6^. In HBV-positive cells, the enrichment of histone ubiquitination (uH2B) and HR factors including RPA, RAD51 and EXO1 at DNA double strand breaks (DSBs) is compromised. This correlates with reduced long-range resection and is largely attributed to ablation of CRL4^WDR70^ activity by HBx. However, the mechanistic role of CRL4^WDR70^ in DNA repair remains undetermined.

Mis-regulation of DSB repair compromises chromosomal stability^7^ and is often characterized by altered usage of non-homologous end joining (NHEJ) and HR. DNA end processing by 5’-to-3’ resection governs HR commitment by generating RPA-coated single-stranded DNA (ssDNA) that subsequently loads Rad51 to form a filament that enables homology search^8^. The cell cycle position and the chromatin context surrounding the DSB site influences ssDNA production^9,10^. DSBs occurring in G1 are repaired by NHEJ while those formed during DNA replication are repaired by HR using the sister chromatid. DSBs occurring post replication are repaired by either NHEJ and HR. The choice of pathway is regulated by the competitive occupancy of 53BP1 and BRCA1^11-13^. A new therapeutic strategy, synthetic lethality (SL), has recently been introduced for cancer subtype-specific chemotherapy and this was first exemplified by the treatment of BRCA-deficient breast cancers with PARP inhibitors (PARPi), including Olaparib and Talazoparib^14^.

DNA repair is also regulated by the 19S regulatory particles (RP), a constituent of the 26S proteasome which degrades ubiquitin-tagged proteins. Distinct from the protease activity sequestered in 20S core particle (CP), the canonical 19S RP recognizes ubiquitinated targets, deubiquitinates and positions them for translocation and unfolding to allow degradation by the CP^15^. The 19S RP is subdivided into a “base” that is constituted of six paralogous AAA+ ATPases (PSMC1-6) plus several non-ATPase proteins (PSMD1^Rpn2^, PSMD2^Rpn1^ and ADRM1^Rpn13^)^16,17^, and a “lid” containing PCI domain proteins (PSMD3, 6, 8, 11-13), MPN domain proteins (PSMD7^Rpn8^ and POH1^Rpn11^), plus the non-PCI/MPN domain subunits including DSS1 (also known as Rpn15 or Sem1). The base and lid are conformationally dynamic and together bind a further subunit, the ubiquitin receptor PSM4^Rpn10^.

As introduced above, in addition to regulating proteolysis by the CP, the RP also performs non-proteolytic roles in the context of chromatin. This was originally identified from the recruitment of a subset of RP proteins to the GAL1-10 promoter, implying a direct role in RNA polymerase II transcription^18^. In the context of DNA repair, the 19S has subsequently been shown to modulate the efficiency of both DSB and nucleotide excision repair^19-21^. In response to DSBs, DSS1^Sem1^ and POH1^Rpn11^ have been shown locate to DSBs sites along with other RP components and regulate Rad52/Pol4, or 53BP1/RAP80 repositioning from DSBs, respectively^22-25^. The number and diversity of 19S associating proteins and functions have obscured the elucidation of its mechanism in chromatin biology and a comprehensive model depicting its interplay with DNA repair machinery at DSBs is lacking.

Here we provide evidence that CRL4^WDR70^ forms a specific complex with the break-associated 19S proteasome (subsequently referred to as CDW19S; <u>C</u>ULLIN4A-<u>D</u>DB1-<u>W</u>DR70-<u>19S</u>) that favours end processing and thus HR. We show that the 19S RP controls both MRE11 and EXO1 nucleases and that CRL4^WDR70^ engages with an EXO1-specific module of the RP and promotes the degradation of ADMR1^Rpn13^. We show that the HBx oncoprotein of HBV, by disrupting the interaction between WDR70 and the Cullin 4 - DDB1 scaffold, disrupts the CDW19S complex leaving scaffold-free WDR70-19S on damaged DNA. As such HBx retards the clearance of ADRM1^Rpn13^, a 19S-associated ubiquitin receptor that we identify as a resection barrier factor. Mechanistically we further show that ADRM1^Rpn13^ is directly ubiquitinylated by CRL4^WDR70^ and that a single lysine mutation in ADMR1^Rpn13^ phenocopies both *WDR70* loss and *HBx* expression. To establish if the consequent HRD can be exploited to treat HBV positive cancer, we examined the impact on tumour-promotion of *in vivo* ablation of murine *Wdr70* with *Trp53* positive and negative backgrounds and demonstrate *Wdr70* knockdown synergises with p53^-/-^ to promote tumorigenesis. Finally, we use cell and patient derived xenograft models to demonstrate that PARP inhibition or systemic knockdown of *ENDOD1*, an atypical nuclease recently shown to be synthetic lethal with HRD^26^, restrained tumour growth. Our data conceptualize SL regimes for HBVHCC based on sensitivity to HRD-targeting therapy.

## Results

### HBx disturbs the balanced choice of DSB repair mechanisms

To assess the impacts of HBV/HBx on DSB repair, we used an I-*Sce*I-induced DSB system to measure sister chromatid repair (HR), single strand annealing (SSA) and NHEJ efficiencies^27^. T43 hepatocytes, which harbour integrated HBV genomes^5^, displayed a modest decrease in HR and SSA relative to the parental HBV-free L02 cells (Fig. 1a). To establish if this HBV-induced HR deficiency can be attributed, at least in part, to HBx ablating CRL4^WDR70^ we ectopically expressed HBx (*HBx*^EE^) in HEK293T cells and in *WDR70*-deleted (*70KO*) derivatives (Fig. 1b). HBx^EE^ biased repair away from HR/SSA and towards NHEJ as expected. *WDR70* deletion did not affect the repair profile beyond that seen when *HBx* is expressed in parental 293T cells, consistent with an epistatic relationship. Thus, HBx induces HRD in part through CRL4^WDR70^ ablation and unbalanced pathway choice.

**Fig. 1.**
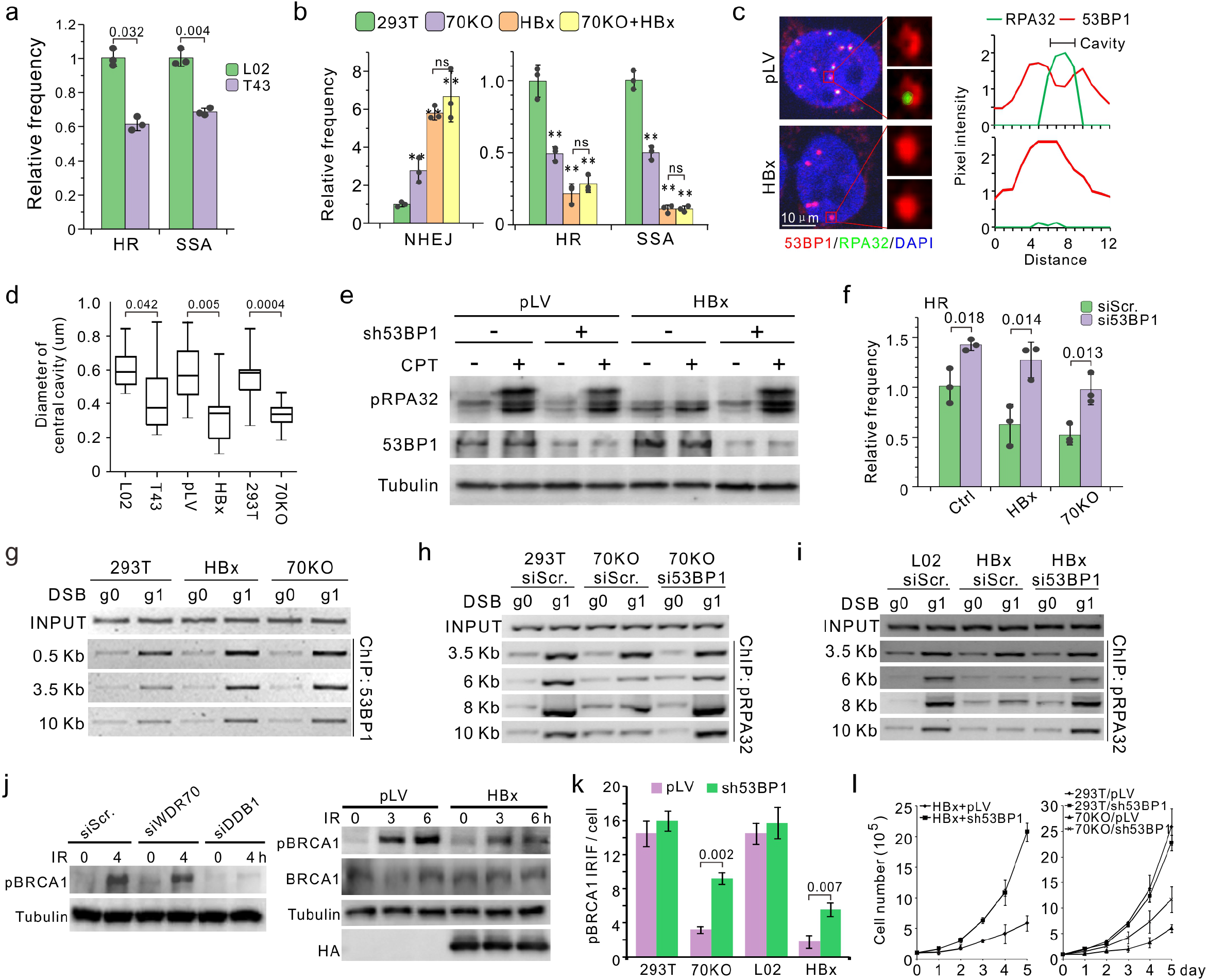
HBx locks DSB-bound 53BP1. (**a**) Repair efficiencies by HR and NHEJ at an I*-Sce*I-induced DSB in HBV carrier cells (T43) were measured by real-time PCR relative to the parental cell line (L02). (**b**) Relative frequency of indicated repair pathways in indicated cells. *p* values by *t*-test for indicated groups are shown. ns: no significant difference. n = 3 biological repeats. Error bars = s.d. (**c**) Left: example confocal images showing 53BP1 (red) and RPA32 (green) IRIF in *HBx-*expressing L02 cells eight hours post-IR and pre-extracted with 0.1% Triton-X100. Right: pixel intensity (vertical) across the maximal central line of individual IRIFs. Precipitation of red line (53BP1) and rising of green line (RPA32) along the vertical axis indicates the central cavity. (**d**) Range of 53BP1 cavity sizes (μm) in indicated cells measured as in (c). n = 3 biological repeats and 25 cavities were included for each sample. box: median and interquartile range; whiskers: minimum and maximum. (**e**) Immunoblot detection of pRPA32 after CPT treatment in *HBx*^EE^ L02 cells expressing, or not, sh*53BP1*. (**f**) Relative HR efficiency for L02 cells pre-treated with *HBx* or si*WDR70* subjected to simultaneous si*53BP1* or control (si*Scr*.) treatments. n = 3 biological repeats. Error bars = s.d. *p* values by *t*-test are shown. (**g**) ChIP assays depicting 53BP1 chromatin loading at indicated distance from the DSB upon expression of gRNA (g1) targeting the *PPP1R12C/p84* locus. g0: control gRNA. (**h**-**i**) ChIP assay of pRPA32 at the indicated distances from the gRNA (g1)-dependent DSB in *70KO* or 293T cells (h) and L02 or L02 *HBx*^EE^ (i) cells with or without *53BP1* knockdown. (**j**-**k**) Immunoblotting (j) and quantification of IRIF (k) for pBRCA1 in L02 cells with the indicated treatments. n = 3 biological repeats. Error bars = s.d. *t*-test. (**l**) Proliferating curve of *HBx*^EE^ or *WDR70* knockout cells with sh*53BP1* expression or not. 3 biological repeats. Error bars = s.d.

53BP1 forms ionizing radiation-induced foci (IRIF) and is required to establish a resection barrier at DSB ends. To activate HR, 53BP1 is subsequently displaced from the focal centre 4 - 8 hours post-IR in a BRCA1-dependent manner^28^. This 53BP1 loss correlates with the accumulation of the ssDNA binding protein RPA within the central cavity of the focus, a process that is dependent on a functional resection machinery. Upon HBV infection, *HBx*^EE^ or *WDR70* loss, a reduction in the size of the central cavity in 53BP1 IRIF that is positive for RPA32 was observed (Fig. 1c,d). Consistent with this si*DDB1* (an obligate partner of all CRL4 complexes), si*WDR70*, or *HBx*^EE^ in L02 cells all prolonged 53BP1 IRIF (Extended Data Fig. S1a-b). As expected, these 53BP1 foci often appeared compacted, lacked sizable central cavities (c.f. Fig. 1d and Extended Data Fig. 1c), and showed reduced co-staining with RPA32 (Extended Data Fig.1d).

We next examined if the *HBx*-induced HRD could be rescued by removing the *53BP1*-mediated resection barrier. Respective to controls, sh*53BP1* restored camptothecin (CPT)-induced RPA32 phosphorylation (pRPA32) in *HBx*^EE^ cells (Fig. 1e), and similarly restored IRIF of pRPA32 and RAD51 recombinase in *70KO* cells (Extended Data Fig. 1e). These observations also correlated with the alleviation of the decreased HR efficacies observed in the I-*Sce*I reporter assay upon *HBx*^EE^ and si*WDR70* (Fig. 1f). Further, using a chromatin immunoprecipitation (ChIP) assay developed for CRISPR-induced DSBs at the *PPP1R12C/p84* locus^5^, we visualized enhanced loading of 53BP1 in *70KO* and *HBx*^EE^ cells at 0.5 - 10 Kb from the break when compared to their respective controls (Fig. 1g). pRPA32 association in *70KO* and *HBx*^EE^ cells was concomitantly restricted 6-10 Kb from the break site and this was reversed by si*53BP1* (Fig. 1h,i). Consistent with the antagonistic role for BRCA1 against 53BP1^29^, HBV infection, *HBx*^EE^ or directly compromising CRL4^WDR70^, all diminished chromatin deposition and IRIF formation of pS1524-phosphorylated BRCA1 (Fig. 1j and Extended Data Fig. 1f). Resembling the restoration of pRPA32, sh*53BP1* increased pBRCA1 foci formation in *HBx*^EE^ and *70KO* cells (Fig. 1k). Congruent with improved DSB repair, impaired cell viability in *HBx*^EE^ and *70KO* cells was ameliorated by sh*53BP1* (Fig. 1l). We conclude that HBx influences DSB repair pathway choice by impeding the removal of a 53BP1-dependent resection barrier that is regulated by CRL4^WDR70^ and BRCA1.

### Assembling CDW19S on break-associated chromatin

To establish how CRL4^WDR70^ interplays with the repair machinery, we purified proteins associated with TAP-tagged Wdr70 from fission yeast. We have previously shown that regulation of resection by CRL4^WDR70^ is conserved in fission yeast^6^. In these experiments we deleted the gene encoding Ddb1, the adaptor that links Wdr70 to Pcu4 (Cullin 4 homolog) to disable the ubiquitin ligase activity of CRL4^Wdr70^ and prevent the potential degradation of binding factors. MALDI-TOF mass spectrometry of sliced gel bands identified three categories of proteins (Fig. 2a and Table S1): histone species (Htb1^H2B^, Hht1^H3.1^), ubiquitination factors (note we see low coverage for Pcu4 and abundant ubiquitin) and, unexpectedly, a range of proteins derived from 19S RP of the proteosome. The panel of RP subunits encompasses the main proteasomal RP subcomplexes including PCI-, MPN-, and ATPase-domain containing proteins. Notably, no peptides from the proteolytic 20S core particle were retrieved.

**Fig. 2.**
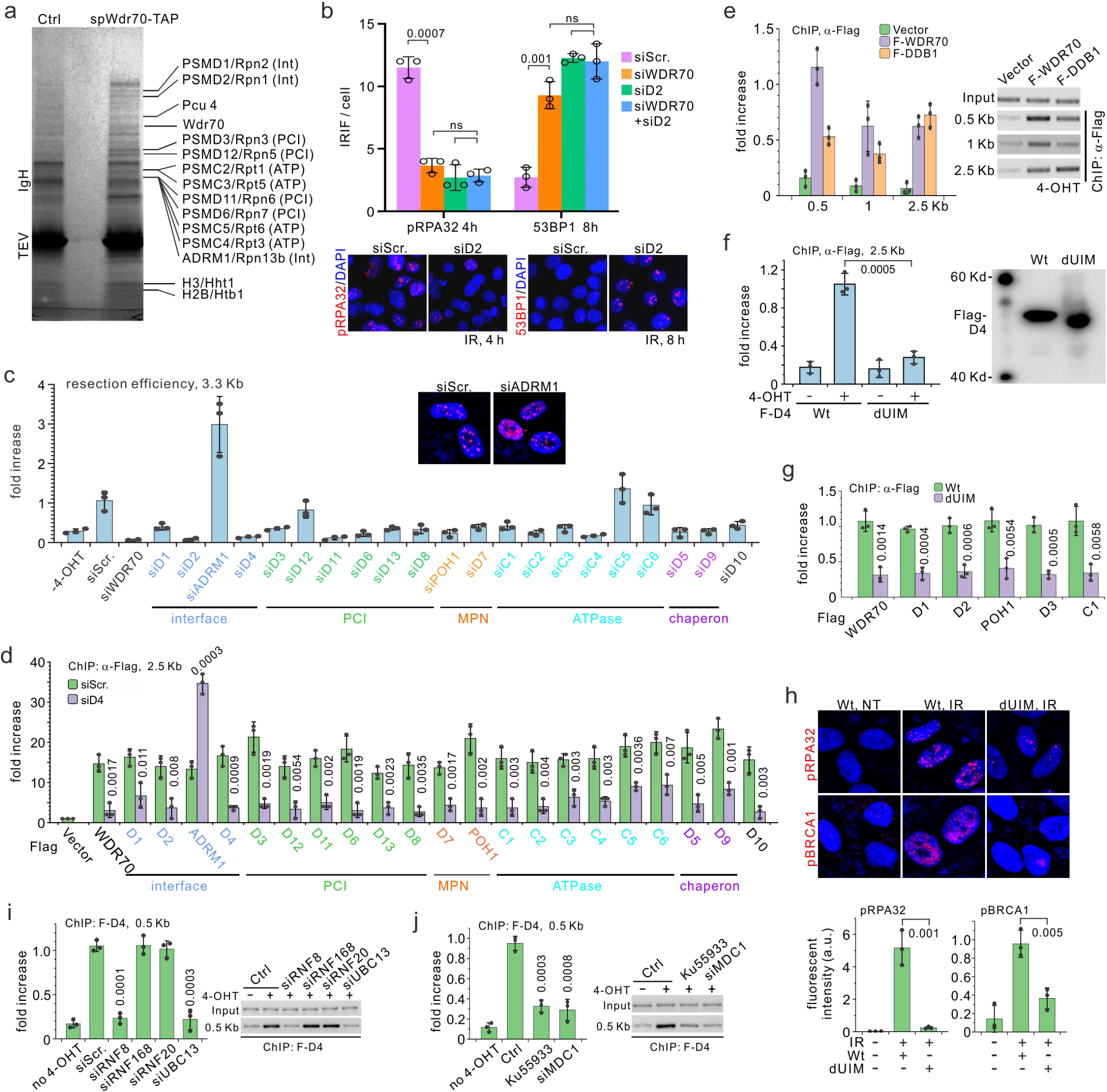
CDW19S engages with DSB proximal chromatin. (**a**) TAP-affinity purified spWdr70-interacting proteins separated by gradient SDS-PAGE. Proteins identified by MALDI-TOF mass spectrometry are shown on the right. See peptide coverage in Table S1. Subdomains of interface (Int), PCI, MPN and ATPase (ATP) are indicated for RP subunits using human and yeast nomenclatures. IgH: heavy chain of rabbit IgG. TEV: Tobacco Etch Virus endopeptidase. (**b**) Representative images and enumeration of pRPA32 and 53BP1 IRIF in HEK293T cells with indicated knockdown treatment. (**c**) Quantification for *Xba*1-based resection assay (See Extended Data Fig. 2f) showing DSB processing in RPE1 cells depleted for the indicated CDW19S subunits. Inset: excessive pRPA32 immunostaining implies hypersection in si*ADRM1*^Rpn13^ RPE1 cells. (**d**) ChIP assay 2.5 Kb distal to an *Asi*SI-induced DSB (DiVA cells) showing break association of the indicated CDW19S subunits upon *PSMD4*^Rpn10^ ablation relative to control siRNA. (**e**) Left: ChIP assay for Flag-tagged WDR70/DDB1 at indicated distances from AsiSI-induced DSB ends. Right: representative PCR products. (**f**) Chip assay of Flag-PSMD4^Rpn10^ 2.5 Kb from an *Asi*SI-induced DSB upon PSMD4 or PSMD4^dUIM^ expression. Immunoblotting for Flag-tagged PSMD4^Rpn10^ is shown on the right panel. (**g**) Equivalent ChIP assay for the indicated Flag-tagged CDW10S subunits in the presence of PSMD4 or PSMD4^dUIM^ expression. (**h**) Top: representative images of pRPA32 and pBRCA1 IRIF in the presence of PSMD4 or PSMD4^dUIM^ expression (4 h post-IR). Nuclei counterstained with DAPI. Bottom: quantification of fluorescent intensity relative to that of wild type. In panels g-h the *PSMD4* and PSMD4^dUIM^ were expressed without Flag tags and are siRNA resistant and cells were co-treated with si*PSMD4*^Rpn10.^.(**i**-**j**) PSMD4^Rpn10^ enrichment at 0.5 Kb from an *Asi*SI-induced DSB quantified by ChIP after treatment with the indicated siRNA or inhibitors. Data normalized to control with 4-OHT induction. All quantification in this figure: n = 3 experimental repeats; error bars: s.d.; *p* values by *t*-test for indicated groups are shown; ns: no statistical significance.

Co-immunoprecipitation assays for WDR70/DDB1 following immunoprecipitation of ectopically expressed Flag-tagged 19S subunits from HEK293T nuclear extracts demonstrated that the CRL4^WDR70^-19S interaction is conserved in human cells (Extended Data Fig. 2a). Consistent with a role in DSB repair, these interactions were enhanced upon CPT treatment (Extended Data Fig. 2b). The engagement of CRL4^WDR70^ to the RP relies on WDR70 since the PSMD4^Rpn10^-DDB1 interaction was compromised in si*WDR70* cells but not *vice versa* (Extended Data Fig. 2c). A functional importance for the interaction is implied by the observation that, as shown previously for *HBx*^EE^ and si*WDR70*^5^, depleting the proteasomal *PSMD2*^Rpn1^ subunit inhibits H2B monoubiquitylation, RPA32 phosphorylation and IRIF (Fig. 2b and Extended Data Fig. 2d). Furthermore, simultaneous silencing *WDR70* and *PSMD2*^Rpn1^ did not exacerbate the pRPA32 IRIF defect, indicative of epistasis between CRL4^WDR70^ and 19S. We thus conclude that CRL4^WDR70^ and the 19S particle form a complex (CDW19S) that influences DSB repair.

To survey the function of the CDW19S complex at DSBs, the DIvA system^30^ (where DNA breaks are generated by ER-tagged *Asi*SI endonuclease upon its nuclear import following 4-OHT treatment) was used to analyse a specific *Asi*SI site on chromosome 1 (89,458,595 - 89,458,603, reference genome hg19) by ChIP. Four hours after 4-OHT treatment, pRPA32, indicative of resection, was observed in control cells between 0.5 and 5 Kb from the break (Extended Data Fig. 2e). Loss of *WDR70* affected distal pRPA32 deposition (2.5 - 5 Kb), whereas proximal (0.5 Kb) resection was not affected. Distal resection (3.3 Kb) was next analysed by digesting genomic DNA with *Xba*I, which is inactive on ssDNA. Uncut ssDNA was quantified by PCR across the digestion site (Extended Data Fig. 2f). Four hours after induction, control cells show evidence of ssDNA at 1 Kb and 3.3 Kb from the break site. To assess the role of CDW19S components in regulating resection, ssDNA was quantified following knockdown of genes encoding CDW19S components or associated proteins. Knockdown of the majority of genes encoding CDW19S subunits, plus those encoding the associated chaperons (PSMD5^Hsm3^ and PSMD9^Nas2^) and PSMD10 proteins, exhibited si*WDR70*-like resection defects (Fig. 2c). *ADRM1*^Rpn13^ depletion stood out, displaying hyperactive ssDNA production relative to control cells. No function for PSMD12^Rpn5^, PSMC5^Rpt6^ and PSMC6^Rpt4^ was observed, despite significant mRNA ablation (Extended Data Fig. 2g).

To examine the recruitment of CDW19S components to break sites, DiVA cells were transformed with plasmids expressing Flag tagged constructs and examined by ChIP following *Asi*SI nuclear import. All RP proteins tested and WDR70/DDB1 were enriched at both 0.5 and 2.5 Kb (Fig. 2d,e and Extended Data Fig. 2h). We conclude that CRL4^WDR70^ decorates 19S RP in a stable CDW19S complex to stimulate resection.

### PSMD4^Rpn10^ recruits CDW19S in an ATM-MDC1-RNF8-dependent manner

19S RP is known to be targeted to chromatin and interplay with DSB-associated ubiquitin conjugates^24,31^. We speculated that the ubiquitin binding directs the RP to the DSBs. PSMD4^Rpn10^, an integral ubiquitin receptor in the RP lid that recruits K63- or K48-linked ubiquitin targets to the proteasome is a promising candidate^32^. Knockdown of *PSMD4*^Rpn10^ abolished DSB recruitment of WDR70 and all RP subunits tested except ADRM1^Rpn13^ (Fig. 2d). In contrast, PSMD4^Rpn10^ was recruited to DSBs irrespective of siRNA treatment for any of the CDW19s subunits tested (Extended Data Fig. 2i). PSMD4^Rpn10^ recognizes ubiquitin chains via a UIM (ubiquitin-interacting motif), mutation of which abrogates ubiquitin affinity^33^. A flag-tagged ectopically expressed UIM-deletion mutant (*PSMD4*^dUIM^) did not associate with DSB (Fig. 2f) and suppressed CDW19S complex assembly (Fig. 2g). Consistent with this PSMD4^dUIM^ expression phenocopied si*WDR70*, repressing resection as monitored by phosphorylated RPA32 and BRCA1 IRIF (Fig. 2h).

To establish if damage signaling was prerequisite for CDW19S DSB association we evaluated PSMD4^Rpn10^ DSB association in conditions that interfere with early signaling events. Silencing the damage-responsive E3 enzyme (RNF8) and its partner E2 (UBC13) effectively abolished PSMD4^Rpn10^ loading 0.5 Kb from the break site (Fig. 2i). Comparable effects were not seen when RNF163 or RNF20 were silenced. RNF8-UBC13 catalyzes K63-polyubiqutination chains, suggesting PSMD4^Rpn10^ is attracted to DSB-associated K63 modified proteins. The RNF8 FHA domain docks to TQxF clusters on MDC1 following their ATM-dependent phosphorylation^34^. Consistent with this, both ATM kinase activity (kinase inhibitor KU55933) and MDC1 (si*MDC1*) were required for DSB recruitment of PSMD4^Rpn10^ (Fig. 2j). We conclude that DSB assembly of CDW19S is initiated in proximity to a DSB by PSMD4^Rpn10^ recognition of RNF8-catalyzed K63 species formed following ATM-MDC1-dependent signaling.

### CDW19S is functionally segregated into MRE11 and EXO1 regulatory modules

To establish if CDW19S has a functional domain structure individual subunits were knocked down and pRPA32 recruitment was assayed 0.5 and 2.5 Kb from an *Asi*SI site by ChIP following *Asi*SI nuclear import. As seen with si*WDR70*, ablation of the majority 19S components led to pRPA32 reduction at 2.5 Kb. For many components, this was effectively reverted by concomitant si*53BP1* treatment (Fig. 3a). The effects of si*PSMD4*^Rpn10^ and si*PSMD5*^Hsm3^ on PRA32 loading was reproduced with a second siRNA and each of these were complemented by respective siRNA resistant plasmids (Extended Data Fig. 3a). Interestingly, resection defects observed upon *PSMD2*^Rpn1^, *PSMD4*^Rpn10^ and MPN (*POH1*^Rpn11^ and *PSMD7*^Rpn8^) depletion were not restored by si*53BP1*. This correlated with a resection defect at 0.5 Kb (Fig. 3b and Extended Data Fig. 3b), implying roles of these four components in initiating proximal resection that does not require CRL^WDR70^ or RP subunits with *WDR70*-like phenotypes.

**Fig. 3.**
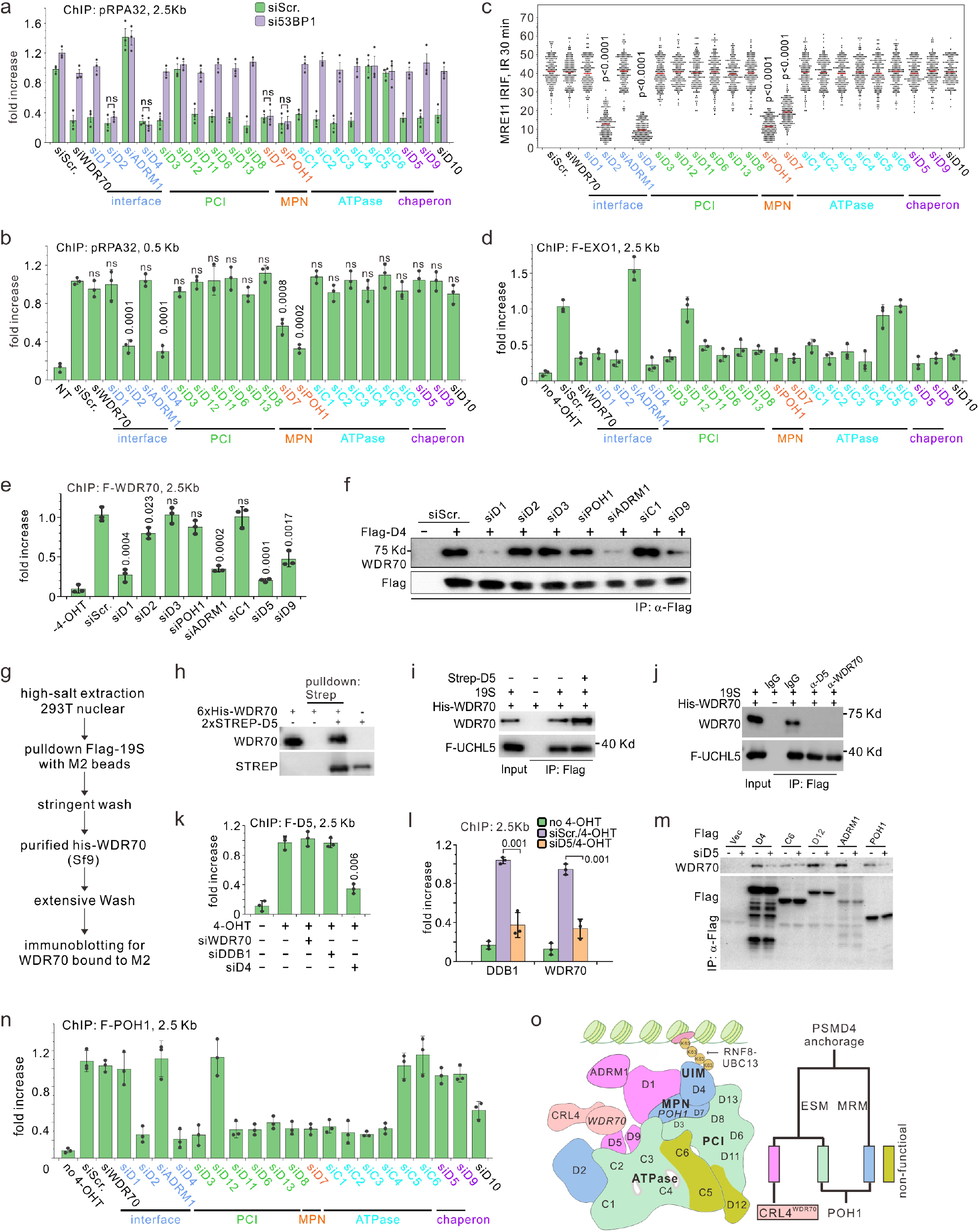
Separate CDW19S modules regulate MRE11 and EXO1 activation. (**a**) ChIP assay showing DSB loading of pRPA32 at 2.5 Kb distal to an *Asi*SI-induced DSB upon silencing of the indicated CDW19S subunits with concomitant siControl or si*53BP1* treatment. Binding efficiencies were normalized to si*Scramble* with 4-OHT induction. (**b**) ChIP assay showing DSB loading of pRPA32 at 0.5 Kb from an *Asi*SI-induced DSB upon silencing of the indicated CDW19S subunits. (**c**) ChIP assay showing loading of EXO1 at 2.5 Kb distal to an *Asi*SI-induced DSB. (**d**) Enumeration for MRE11 foci upon silencing of individual CDW19S subunits in RPE1 cells. IF was carried out 30 min post-IR. n = 3 biological repeats, 50 cells counted for each repeat. Error bars = s.d. *p* values by *t*-test are shown. (**e**) ChIP assay for loading of Flag-tagged WDR70 2.5 Kb distal to an *Asi*SI-induced DSB following siRNA treatment of the indicated proteins. Data normalized to si*Scramble* with 4-OHT induction. (**f**) co-IP of PSMD4^Rpn10^ and WDR70 from chromatin fractions of CTP-treated HEK293T cells with or without ablating indicated RP components. (**g**) Schematic showing the high-salt procedure for screening 19S components for direct engagement with WDR70. (**h**) Pull-down assay using purified WDR70 and PSMD5^Hsm3^. (**i**-**j**) *In vitro* pull-down assay for purified WDR70 (0.5 µg) and 19S proteasome (2 µg), the latter containing Flag-UCHL5. Recombinant PSMD5^Hsm3^ (1 µg, i) or specific antibodies (0.5 µg, j) were added into the reaction. (**k**-**l**) ChIP assay for loading of Flag-PSMD5^Hsm3^ (k) or Flag-DDB1/WDR70 (l) 2.5 Kb distal to an *Asi*SI-induced DSB following indicated siRNA treatments. (**m**) Immunoprecipitation as in (f) for endogenous WDR70 and indicated Flag-tagged 19S subunits with or without *PSMD5*^Hsm3^ silencing. (**n**) ChIP assay showing DSB loading of POH1^Rpn11^ at 2.5 Kb distal to the an *Asi*SI-induced DSB following siRNA treatment against the indicated CDW19S subunits. (**o**) Illustration for MRM and ESM modules of CDW19S. The architecture of 19S is based on Lauren Budenholzer’s description^15^. Unless otherwise indicated, all quantification in this figure: n = 3 experimental repeats; error bars: s.d.; *p* values by *t*-test are shown for indicated groups; ns: no statistical significance.

To further explore these distinctions, we analysed the formation of MRE11 IRIF. CRL4^WDR70^ depletion or *HBx*^EE^ does not diminish the proximal nuclease (MRE11) from DNA ends (ref^5^). However, the ablation of *PSMD2*^Rpn1^, *PSMD4*^Rpn10^, PSMD7^Rpn8^ or POH1^Rpn11^ prevented the formation of MRE11 IRIF 30 minutes and 2 hours post IR (Fig. 3c and Extended Data Fig. 3c,d). This correlates with their requirement for both proximal (0.5 Kb) and distal (2.5 Kb) DNA processing (see Fig. 3a,b). We therefore categorize an MRE11-regulatory module (MRM) within CDW19S that is necessary to initiate end processing.

We next analysed chromatin association of the long-range exonuclease EXO1. Distal EXO1 association (2.5 Kb) was reduced upon silencing WDR70 and CDW19S subunits that phenocopy *WDR70*, but the same impact was not observed upon si*ADRM1*^Rpn13^, si*PSMD12*^Rpn5^, si*PSMC5*^Rpt6^ or si*PSMC6*^Rpt4^ (Fig. 3d). Thus, we conclude that CRL4^WDR70^, together with a majority of RP components, constitutes an EXO1-specific module (ESM) that regulates extensive resection. Again, we observed that depletion of ADRM1^Rpn13^ stands out as promoting increased distal EXO1 and pRPA32 recruitment when compared to control cells suggesting it limits, rather than promotes, distal resection (c.f. Fig. 2c, Fig. 3a,d, Fig. 3d). Since PSMD4^Rpn10^ is required for CDW19S ubiquitin association and CDW19S assembly at DSBs, these data reaffirm that PSMD4^Rpn10^ is required to launch CDW19S assembly and suggest that two separate modular functions, MRM and ESM promote proximal and distal resection, respectively, by regulating distinct nuclease activities.

### PSMD1^Rpn2^-ADRM1^Rpn13^-PSMD5^Hsm3^-PSMD9^Nas2^ are required for CRL4^WDR70^ recruitment

To establish how CRL4^WDR70^ docks to the RP, WDR70 ChIP was exploited in the DiVA system combined with targeted siRNA. CRL4^WDR70^ loading at 2.5 Kb following *Asi*SI nuclear import was significantly compromised by PSMD1^Rpn2^, ADRM1^Rpn13^, PSMD5^Hsm3^ and PSMD9^Nas2^ ablation (Fig. 3e), implying an interface dependent on these four ESM subunits tethers CRL4^WDR70^ to the RP. Indeed, co-immunoprecipitation between WDR70 and Flag-PSMD4^Rpn10^ (the recruiting factor for CDW19S) in chromatin fractions was abolished in the absence of PSMD1^Rpn2^, ADRM1^Rpn13^, and PSMD9^Nas2^ but was maintained when representative interface (*PSMD2*^Rpn1^), PCI (*PSMD3*^Rpn3^), MPN (*POH1*^Rpn11^) or ATPase (*PSMC1*^Rpt2^) subunits were depleted (Fig. 3f).

To attempt to identify a direct docking site of CRL4^WDR70^, 19S particles were dissociated in nuclear HEK293T extracts using high-salt buffer (600 mM NaCl) and incubated with His-WDR70 (112-654 aa) purified from Sf9 cells (Fig. 3g).The sole putative WDR70 binding partner identified was PSMD5^Hsm3^, a chaperon that contributes to RP assembly and only loosely associates with mature 19S^35^. This interaction was confirmed by co-precipitation of recombinant His-WDR70 and Strep-PSMD5^Hsm3^ (Fig. 3h and Extended Data Fig. 3e). Recombinant WDR70 also displayed affinity with commercial purified RP (R&D, E-367) (Fig. 3i). This WDR70-19S interaction was attributed to low amount of PSMD5^Hsm3^ that co-purified with 19S (Extended Data Fig. 3f,g), since it was boosted by adding additional (0.5 ug) recombinant PSMD5^Hsm3^ and was weakened by pre-treating with specific antibodies against either PSMD5^Hsm3^ or WDR70 (Fig. 3i,j). Consistent with other CDW19S subunits, PSMD5^Hsm3^ deposition at DSB sites required PSMD4^Rpn10^ but not WDR70 or DDB1 (Fig. 3k) and si*PSMD5*^Hsm3^ impeded the DSB association of both DDB1 and WDR70 (Fig. 3l). si*PSMD5*^Hsm3^ also attenuated co-immunoprecipitation of endogenous WDR70 with Flag-PSMD4^Rpn10^, PSMD12^Rpn5^, ADRM1^Rpn13^ and POH1^Rpn11^ (Fig. 3m). PSMC6^Rpt4^ was mildly affected. We posit that PSMD5^Hsm3^, together with ADRM1^Rpn13^, PSMD1^Rpn2^ and PSMD9^Nas2^ chaperones CRL4^WDR70^ to the RP, reminiscent of the contribution of chaperons to the stepwise assembly of RP base subcomplexes^35^.

Previous reports demonstrate that POH1^Rpn11^ regulates DSB repair by inhibiting NHEJ while promoting HR^23,24^. Consistent with this, we observed increased proximal (0.5 Kb) KU80 binding in the DiVA system upon si*POH1*^Rpn11^. Concomitant depletion of both POH1 and MRE11 did not lead to a further increase in KU80 binding (Extended Data Fig. 3h), implying that POH1^Rpn11^ loss leads to MRE11 destabilization. si*POH1*^Rpn11^ also resulted in increased 53BP1 levels at later timepoints (8h) following IR treatment (Extended Data Fig. 3i) but did not impact 53BP1 recruitment at 30 min post-IR. This would be consistent with an additional role for POH1^Rpn11^ in distal resection as part of the ESM. In support of this, ChIP analysis using the DiVA system demonstrated the presence of POH1^Rpn11^ at 2.5 Kb (Fig. 3n). Notably, ablation of the majority of CDW19S subunits, but not WDR70, ADRM1^Rpn13^, the associated (PSMD5^Hsm3^, PSMD9^Nas2^ and PSMD10), or non-functional components (PSMD12^Rpn5^, PSMC5^Rpt6^, PSMC6^Rpt4^), affected POH1^Rpn11^ deposition. This indicates that distinct 19S components specialize in recruiting distinct enzymatic activities. Collectively, these data distinguish at least two architecturally separable subdomains within ESM that respectively sustain the functions of distal POH1^Rpn11^ and CRL4^WDR70^. They act in parallel to overcome the 53BP1-dependent resection barrier at DSBs (Fig. 3o).

### CRL4^WDR70^ regulates the ubiquitin-dependent degradation of break-associated ADRM1^Rpn13^

CRL4^WDR70^ is a Ub ligase that promotes resection, predicting that it targets an anti-resection factor for degradation and implicating such a factor to be more abundant on broken chromatin in the absence of CDW19S. ADRM1^Rpn13^ is highly associated with chromatin in the absence of CDW19S (c.f. Figure 2d) and its ablation results in increased resection (c.f. Fig 2c) and RPA/EXO1 loading (c.f. Figure 3a,c). We therefore tested ADRM1^Rpn13^ recruitment to an *Asi*SI-induced DSB by ChIP, with or without siRNA of *WDR70*. ADRM1^Rpn13^ was recruited more abundantly upon si*WDR70*, particularly at the distal (2.5 Kb) region (Extended Data Fig. 4a), and this was accompanied by increased EXO1 recruitment (Extended data Fig4b; left). Importantly, co-depletion of *ARDM1*^Rpn13^ and *WDR70* restored EXO1 recruitment at 2.5 Kb when compared to *WDR70* depletion alone (Extended Data Fig. 4b; right).

Like PSMD4^Rpn10^, ADRM1^Rpn13^ is an RP ubiquitin receptor^36^. Interestingly, DSB recruitment of ADRM1^Rpn13^ is not dependent on PSMD4^Rpn10^ (cf. Fig. 2d) and can therefore occur independently of CDW19S. However, ADRM1^Rpn13^ influences the loading of WDR70 as part of the recruiting platform dependent on PSMD1^Rpn2^, ADRM1^Rpn13^, PSMD5^Hsm3^ and PSMD9^Nas2^ (c.f. Fig. 3e). ADRM1^Rpn13^ encodes a C-terminal DEUBAD domain (265-407 aa) and a conserved N-terminal pleckstrin-like receptor for ubiquitin (Pru) (1-150 aa, ^36^) that preferably binds to K48 polyubiquitin chains via a triple-residue motif. Substitution of *I75R, F76R*, and *D79N* (ADRM1^mIFD^) abolishes the ubiquitin affinity (Extended Data Fig. 4c). Expressing a truncation of the Pru-domain (Extended Data Fig 4d), but not truncation of the DEUBAD domain, reduced enrichment in CPT-treated chromatin, while expressing ADRM1^mIFD^ (Extended Data Fig 4e) reduced enrichment at an AsiSI-dependent DSBs.

We next expressed siRNA resistant ADRM1^Rpn13^ or ADRM1^mIFD^ in cells treated with concomitant si*ADRM1*^Rpn13^ and si*WDR70*. si*ADRM1*^Rpn13^ restores resection in the absence of *WDR70*, as measured by ChIP of RPA. siRNA-resistant wild type ADRM1^Rpn13^ reversed this, whereas siRNA-resistant ARDM1^mIDF^ did not (Fig. 4a), suggesting that Ub-association via the Pru domain is required for the anti-resection function. To investigate the relationship between ADRM1^Rpn13^ and 53BP1, a resection barrier protein, we examined the intensity of 53BP1 IRIF (Fig. 4b). In the absence of ADRM1^Rpn13^, 53BP1 foci intensity is reduced. As shown previously, si*WDR70* increases 53BP1 foci intensity, and this is reversed by concomitant si*ADRM1*^Rpn13^, suggesting the ADRM1^Rpn13^ acts as the upstream barrier protein. Therefore, CRL4^WDR70^ counteracts ADRM1^Rpn13^ and loss of ADRM1^Rpn13^ function obviates the need for CRL4^WDR70^ to ^5^promote extensive resection.

**Fig. 4.**
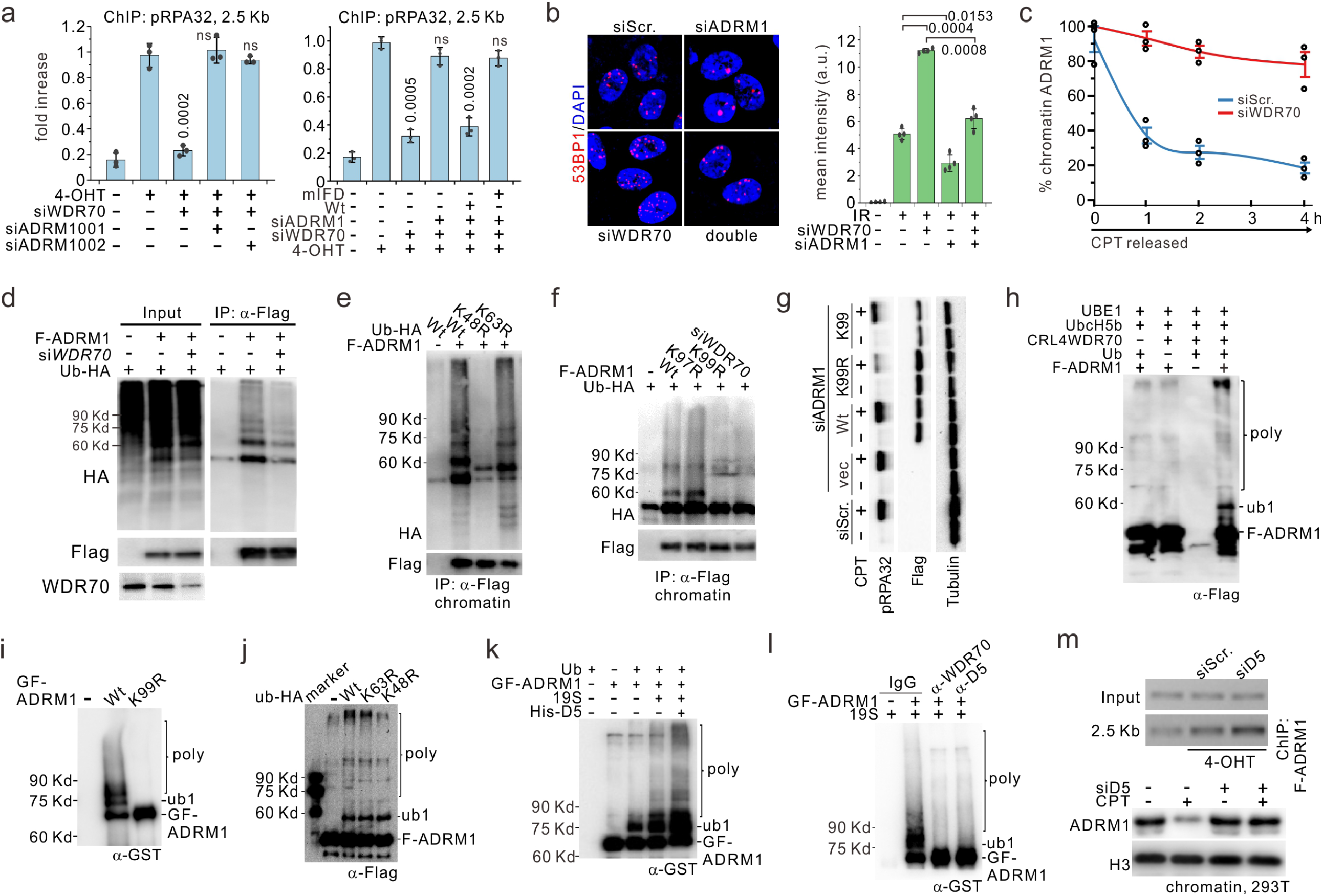
CRL4^WDR70^ targets ADRM1^Rpn13^ for UPS degradation. (**a**) left: ChIP assay showing DSB loading of pRPA32 at 2.5 Kb distal to the an *Asi*SI-induced DSB upon silencing *ADRM1*^Rpn13^ (two different siRNAs) or WDR70. Right: equivalent assay expressing si001-resistant Wt or m*IFD* mutant *ADRM1*^Rpn13^. (**b**) Representative images (left) and fluorescent quantification (right) for 53BP1 IRIF (8 h post-IR) in the presence or not of si*ADRM1*^Rpn13^, with or without simultaneous silencing of *WDR70*. (**c**) Protein abundance of ADRM1^Rpn13^ in chromatin fraction as measured by immunoblotting. Cells were treated with 2 µM CPT for 1 hours and released into drug-free medium. (**d**-**f**) Ub pulldown assay to identify polyubiquitinated species of Flag-ADRM1^Rpn13^ upon treatment with si*WDR70* (d), expression of ubiquitin variants (e) or ADRM1^Rpn13^ K>R mutants (f). Cells were pre-treated with 2 µM CPT for 2 hours. (**g**) Immunoblotting for CPT-induced pRPA32 upon expressing Wt or indicated K>R mutant of Flag-ADRM1^Rpn13^ that are siADRM1-001 resistant. (**h**) Reconstitution of ADRM1^Rpn13^ ubiquitination catalysed with purified proteins. F-ADRM1: Flag-tagged ADRM1. (**i**) Equivalent full reaction using Wt or *K99R* versions. GF-ADRM1: GST-Flag-tagged ADRM1. (**j**) Equivalent full reaction substituting Wt ubiquitin with *K48R* or *K63R* proteins. (**k**-**l**) ADRM1^Rpn13^ ubiquitination reconstituted with addition of purified 19S and His-PSMD5^HSM3^ (k), or in the presence of α-WDR70 or α-PSMD5^HSM3^ (l). (**m**) ChIP assay for DSB-associated (2.5 kb from an *Asi*SI-induced DSB; upper panel) and chromatin-bound ADRM1^Rpn13^ following CPT treatment with or without *PSMD5*^HSM3^ silencing. All quantification in this figure: n = 3 experimental repeats; error bars: s.d.; *p* values by *t*-test are shown for indicated groups; ns: no statistical significance.

Our data suggest that ADRM1^Rpn13^ degradation by the E3 ligase activity of CRL4^WDR70^ and UPS system is required for distal resection. To further test this we explored the stability of chromatin-associated ADRM1^Rpn13^ upon CPT treatment (Fig. 4c and Extended Data Fig. 4f). ADRM1^Rpn13^ is depleted from the chromatin fraction of CPT-challenged 293T cells, but remained stable upon co-treatment with si*WDR70* or proteasomal inhibitor (MG132). Thus, damage-induced ADRM1^Rpn13^ protein turnover is WDR70 and UPS-dependent. Furthermore, polyubiquitination of chromatin-associated ADRM1^Rpn13^ is decreased upon si*WDR70* (Fig. 4d) and impaired by the expression of a *K48R*, but not *K63R*, ubiquitin mutant (Fig. 4e).

To establish where chromatin-bound ADRM1^Rpn13^ is ubiquitinated we identified seven conserved lysine residues (Extended Data Fig. 4g) between human and *S. pombe* ADRM1^Rpn13^ (*S. pombe* has two homologs; Rpn13a and Rpn13b). Expressing individual Flag Tagged K>R mutations in the presence of HA-tagged ubiquitin and analyzing the Ub profile identified that mutation of *K99* (ADRM1^K99R^), but not other lysine residues (i.e. K97), abrogated Ub-conjugated ADRM1^Rpn13^ species from the chromatin fraction (Fig. 4f). Strikingly an ADRM1^K99-only^ mutant was sufficient for ADRM1^Rpn13^ polyubiquitination and this remained *WDR70*-dependent (Extended Data Fig. 4h). Consistent with K99 determining ADRM1^Rpn13^ regulation, ADRM1^K99R^ was associated more robustly at 2.5 Kb distal to an *Asi*SI site when compared wild type and *K99*-only ADRM1^Rpn13^ and this was epistatic with *WDR70* loss (Extended Data Fig. 4i). This reduced ADRM1^K99R^ ubiquitination and resulting increase in DSB association correlated with reduced pRPA32 after CPT treatment when si*ADRM1*^Rpn13^ was complemented with siRNA-resistant ADRM1^K99R^ relative to that seen upon complementation by expressing either wild type or ADRM1^K99-only^ (Fig. 4g). Collectively, these results support the hypothesis that CRL4^WDR70^ promotes the ubiquitination and degradation of chromatin-bound ADRM1^Rpn13^ and that compromising this modification stalls distal resection.

### 19S complex boosts ADRM1^Rpn13^ ubiquitination via CRL4^WDR70^-PSMD5^HSM3^ engagement

To corroborate the direct ubiquitination of ADRM1^Rpn13^ by CRL4^WDR70^, and explore the role of the 19S RP, CRL4^WDR70^ subunits (His-tagged CULLIN4A, DDB1, RBX and truncated WDR70^112-654^) were co-expressed in sf9 insect cells to assemble the E3 ligase (Extended Data Fig 4j). CULLIN4A-DDB1-RBX-WDR70^112-654^ (CRL4^WDR70^) was supplemented with a ubiquitin reconstitution system (activating enzyme (E1), conjugation factor (E2, UbcH5b) and biotinylated ubiquitin) and tested for in vitro ubiquitinylation of bacterially expressed Flag-ADRM1^Rpn13^ (Extended Data Fig 4k). Strong ubiquitin conjugation activity was observed (Fig. 4h) with signals of mono- and poly-ubiquitination which were absent when the reaction was conducted with ADRM1^K99R^ (Fig. 4i). Consistent with the *in vivo* ubiquitination of ADRM1^Rpn13^, conjugates are K48-linkage (Fig. 4j) and are catalyzed by UbcH5b, but not UbcH5a or UbcH6 (Extended Data Fig 4l).

In vivo, an interface dependent on PSMD1^Rpn2^, ADRM1^Rpn13^, PSMD5^Hsm3^ and PSMD9^Nas2^ RP subunits attracts CRL4^WDR70^ to DSBs, juxtaposing the E3 ligase activity to substrates (i.e. ADRM1^Rpn13^). We thus examined how purified 19S affected CRL4^WDR70^-dependent ADRM1^Rpn13^ ubiquitination. When purified 19S complex was supplemented into the CRL4^WDR70^-ADRM1^Rpn13^ (GST-tagged) reconstitution reaction ubiquitination activity was significantly stimulated. This was further boosted by concomitant addition of recombinant His-PSMD5^Hsm3^ (Fig. 4k) and could be inhibited with by including α-D5 or α-WDR70 antibodies (Fig. 4l). Consistent with this, *in vivo* si*PSMD5*^Hsm3^ resulted in accumulation of ADRM1^Rpn13^ at an *Asi*SI-induced DSB and stabilization after CPT treatment (Fig. 4m). Thus, CRL4^WDR70^ acts as an RP-associated E3 ligase to catalyze the ubiquitination and removal of ADRM1^Rpn13^ from DSB associated chromatin, thus releasing a barrier to long-range resection.

### HBx attenuates the ESM to generate a viral HRD subtype

HBx disintegrates the CRL4^WDR70^ complex by displacing WDR70 from CUL4A-DDB1 via an H-box (88-100 aa)^5,37^. To investigate the impact of HBx on CDW19S, CRL4^WDR70^ was co-immunoprecipitated with Flag-tagged PSMD2^Rpn1^. In L02 cells, the RP interacted with both DDB1 and WDR70. In HBV+ T43 cells, PSMD2^Rpn1^ maintained its association with WDR70 while partially dissociating from DDB1 (Fig. 5a). Treatment with si*HBx* showed this decreased association to be HBx-dependent (Extended Data Fig. 5a). A similar experiment in *HBx*^EE^ L02 cells confirmed that HBx expression compromised the DDB1, but not the WDR70, interaction with PSMD2^Rpn1^ (Fig. 5b). Given that DDB1 association with damaged chromatin is WDR70-dependent^6^, we reasoned that HBx detaches the CRL4-DDB1 scaffold from ESM, leaving partially assembled CDW19S (WDR70-19S) at breaks. Indeed, expressing *HBx* in DIvA cells significantly reduced DDB1 loading at *Asi*SI-induced DSBs without affecting other CDWS19 components (Fig. 5c). Thus, HBx disrupts ESM and leaves ‘torso’ CDW19S complexes at DSBs.

**Fig. 5.**
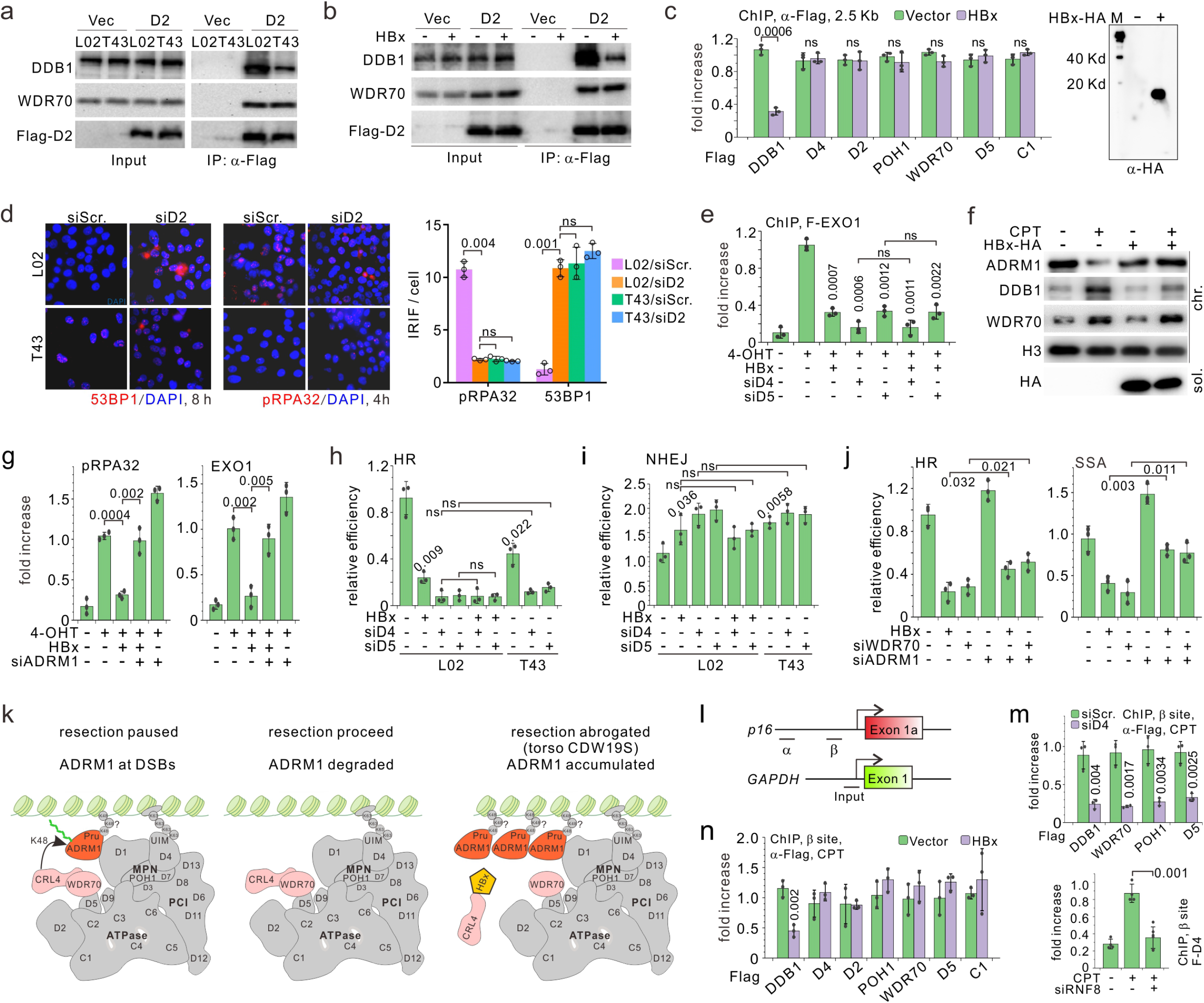
Viral HRD caused by compromised CDW19S by HBx. (**a**-**b**) Immunoprecipitation of DDB1-WDR70 by Flag-tagged PSMD2^Rpn1^ from L02 or T43 cell extracts (a), or in L02 cells expressing, or not, HBx (b). (**c**) ChIP assay for the Flag-tagged CDW19S subunits 2.5 Kb from a *Asi*SI-induced DSB in the presence, or not, of *HBx* expression. ChIP (DIvA cells all with 4-OHT induction) was normalized to DDB1 value without *HBx* expression. (**d**) Representative images (left) and quantification (right) of pRPA32 and 53BP1 foci at the indicated hours post-IR in L02 or T43 cells with si*PSMD2*^Rpn1^ treatment or control treatment. (**e**) ChIP assay for EXO1 loading 2.5 Kb from an *Asi*SI-induced DSB in the presence of the indicated treatments. Values normalized to control EXO1 with 4-OHT. (**f**) Chromatin and soluble nuclear fractionation of indicated proteins upon CPT insult in the presence, or not, of HBx-HA expression. (**g**) ChIP assay for EXO1 or pRPA32 loading 2.5 Kb from an *Asi*SI-induced DSB in the presence of the indicated treatments. (**h**-**i**) Repair efficacies for HR (h) and NHEJ (i) measured by I-*Sce*I DSB system in L02 and T43 cells with indicated treatments. (**j**) HR/SSA repair assay in the presence of *HBx*^EE^ or si*WDR70* with or without *ADRM1*^Rpn13^ depletion in L02 cells. (**k**) Schematic showing HBx dissociates the CRL4 backbone and leads to ‘torso’ CDW19S on chromatin. Consequently, resection fails to proceed due to excessive deposition of ADRM1^Rpn13^ causing a resection barrier. (**l**-**n**) ChIP assays for the indicated Flag-tagged CDW19S subunits at promoter regions of *p16*^INK4a^ in L02 cells following CPT treatment (0.4 µM, 12 hours), siRNA or *HBx*^EE^ treatments as indicated. Amplicons of β (500 bp upstream transcription starting site) and α (that 1300 bp upstream of β) sites at *p16*^INK4a^ and input amplicon at *GAPDH* promoter are shown (l). All quantification in this figure: n = 3 experimental repeats; error bars: s.d.; *p* values by *t*-test are shown for indicated groups; ns: no statistical significance.

To evaluate the biological outcome of torso-CDW19S, we analyzed repair at CRISPR-induced DSBs in T43 cells. This system recapitulates the mechanistic features surrounding *Asi*SI-induced breaks in that si*PSMD4*^Rpn10^ dissociates WDR70 from the break site, but si*WDR70* does not dissociate PSMD4^Rpn10^ (Extended Data Fig. 5b,c). Both si*PSMD2*^Rpn1^ in L02 cells and HBV+ in T43 cells result in increased 5BP1 IRIF 8 hours after irradiation and decreased pRPA32 IRIF. The combined treatment is not additive (Fig. 5d) indicating that HBx acts through the CDW19S pathway. To establish if HBx influences the MRE11- or EXO1-specific functions of CDW19S we examined the effect of *HBx*^EE^ on EXO1. HBx expression reduced the DSB association of EXO1 at an *Asi*SI-induced break and this was not exacerbated when *PSMD4*^Rpn10^ or *PSMD5*^Hsm3^ concomitantly depleted (Fig. 5e). We have previously reported that HBx does not affect MRE11 kinetics^5^. We next examined the influence of HBx on ADRM1^Rpn13^. *HBx*^EE^ resulted in accumulated chromatin-bound ADRM1^Rpn13^ but decreased DDB1 association upon CPT treatment (Fig. 5f), and decreased ChIP of RPA32 and EXO1 distal (2.5 Kb) to an *Asi*SI-induced DSB (Fig. 5g). Importantly, si*ADRM1*^Rpn13^ reversed the decrease in RPA32 and EXO1 break association. To establish how HBx influences repair pathway choice we analyzed I-*Sce*I-induced repair outcomes. These demonstrate that the imbalanced activities of NHEJ/HR apparent upon HBV+ (T43) or HBx^EE^ are epistatic with siRNA of CDW19S subunits required for either Ub-dependent DSB anchorage (PSMD4^Rpn10^) or specific for the regulation of EXO1 (ESM, PSMD5^Hsm3^) (Fig. 5h,i). As expected, the HBx-dependent defect in pathway choice (HR and SSA) was, in part, reversed by si*ADRM1*^Rpn13^ (Fig. 5j). Thus, torso-CDW19S resulting from HBx induces a viral HRD subtype by inhibiting the ESM module of CWD19S and impeding the removal of ADRM1^Rpn13^ (Fig. 5k) and associated EXO1 resection.

### CDW19S functions in gene expression

A recent report suggested the involvement of CRL4^WDR70^ in regulating gene expression of repair factors (i.e. *BRCA1, BRCA2* and *RAD51*) via the DREAM complex, raising an alternative explanation of its role in DNA repair^38^. Partially mirroring this observation, routine gene silencing in this study to ablate expression of *WDR70, PSMD5*^Hsm3^ or *PSMD4*^Rpn10^ moderately decreased the transcription of the above repair genes but did not have a major influence on associated protein levels (Extended Data Fig. 5d). This contrasts with the clear repair defects we observed, suggesting direct roles for CDW19S in DSB repair. Given the well documented link between non-proteolytic RP activity and gene expression via epigenetic control^39-43^, a direct role in DNA repair is not mutually exclusive with the participation of CDW19S in gene expression. In HCC the expression of *p16*^INK4a^ cancer suppressor is subverted in HBV positive cases^44,45^. We thus examined the induction of CPT-induced mRNA and protein and levels of *p16INK4A* / p16^INK4a^. We observed that HBV infection, HBx^EE^ (Extended Data Fig 5e,f) and si*WDR70* (Extended Data Fig 5g) each attenuated *p16*^INK4a^ expression and HBx status was epistatic with si*WDR70*. Consistent with WDR70 acting via the CDW19S in regulating transcription, depleting the MRM/ESM subunits, with the exception of *ADRM1*^Rpn13^, or depleting *RNF8* phenocopied si*WDR70* regarding *p16*^INK4a^ expression (Extended Data Fig. 5h,i).

By ChIP we observed that DDB1 and WDR70 are increased proximal to the promoter (β site, 500 bp upstream of TSS) relative to the distal α site (1800 bp upstream TSS, Fig. 5l and Extended Data Fig. 5j). This promoter proximal binding relies on the ubiquitin binding anchor subunit of CDW19S (PSMD4^Rpn10^) and RNF8, as was seen for DSB association (Fig. 5m). To establish if, like at DSB sites, HBx induces a torso-CDW19S at the *p16*^INK4a^ promoter, we tested a subset of subunits by ChIP. Only DDB1 was dissociated from the β site HBx expression (Fig. 5n). Finally, we examined the induction of H2B monoubiquitination, which is indirectly influenced by WDR70 loss at sites of DSBs^5,6^. β-associated H2B monoubiquitination was decreased upon si*WDR70*, si*DDB1* or HBx^EE^ (Extended Data Fig. 5k). Thus, RNF8-triggered PSMD4^Rpn10^ association of the RP and its decoration with CRL4^WDR70^ accentuates various RP functions on chromatin.

### Concurrent loss of *TP53* and *WDR70* promotes carcinogenesis

Curtailing CDW19S-mediated regulation of HR could promote hepatocarcinogenesis by inducing genomic instability. In genomically unstable *BRCA1* deficient cells, p53 counteracts tumorigenesis by triggering apoptosis when elevated DNA damage is detected^46^. An direct interplay between HBx and p53 has previously be demonstrated^47^, with HBV/HBx inhibiting the p53-dependent transactivation of death targets. This activity is independent of WDR70 (Extended Data Fig. 6a). We find that HBV+ T43 cells were insensitive to cisplatin and could be sensitised by the p53 activator, nutlin (Extended Data Fig. 6b). Consistent with this, the growth defect caused by *WDR70* depletion was partially alleviated by concomitant *TP53* ablation (Fig. 6a). We propose that p53-dependent apoptosis, which should eliminate damaged cells upon the subversion of CDW19S by HBx, is attenuated by the known suppression of p53 activity by HBx.

**Fig. 6.**
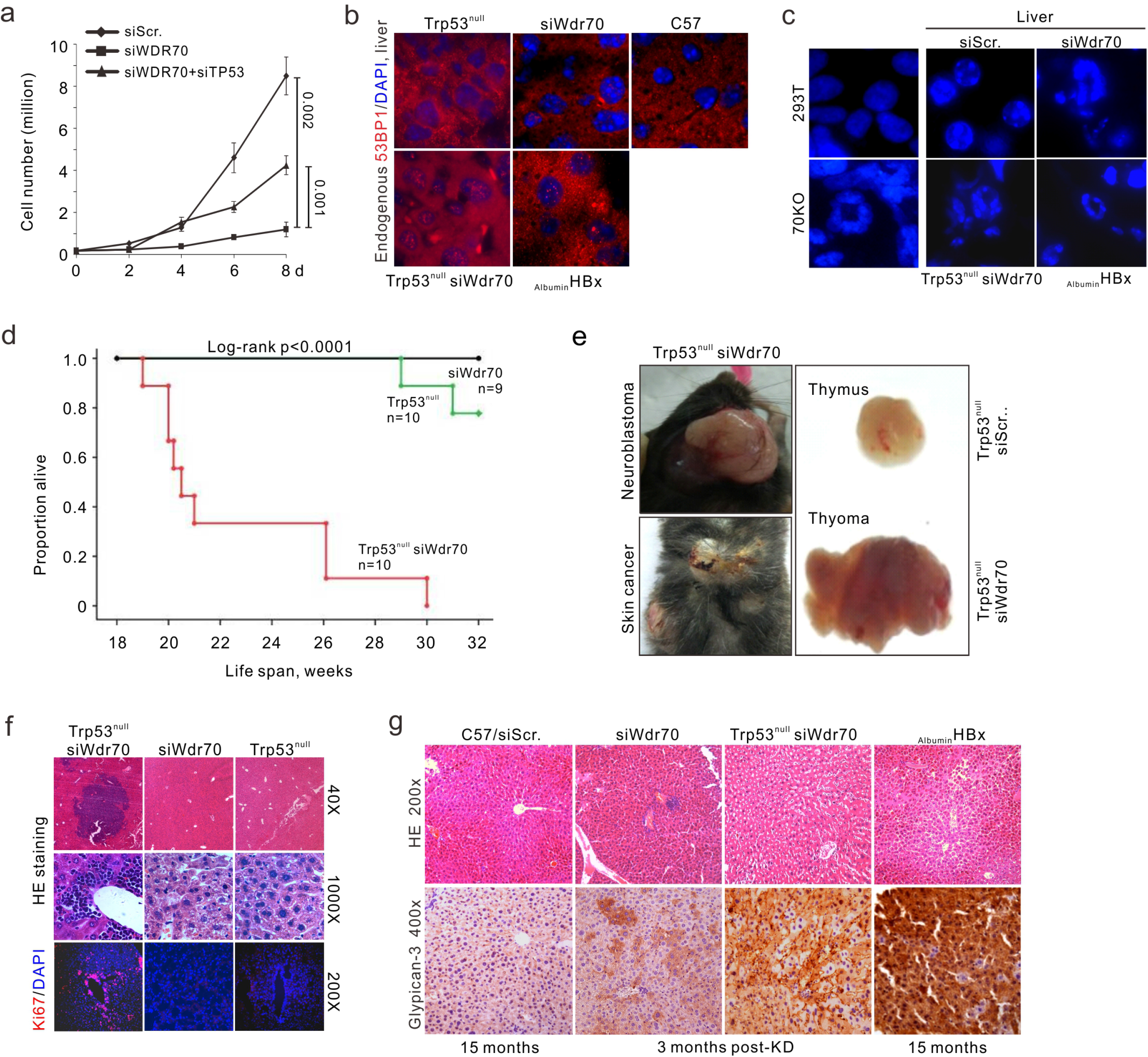
Coincident loss of *TP53* and *WDR70* promotes cancer initiation. (**a**) Growth curve over time following silencing of *WDR70* and/or *TP53* in L02 hepatocytes. n = 3 experimental repeats; error bars: s.d.; *p* values shown for indicated groups; *t*-test. (**b**) Immunostaining for endogenous 53BP1 foci in mouse livers dissected from the indicated genotypes. (**c**) DAPI staining for aberrant nuclear morphology in *70KO* and parental control cells and liver sections from mice of the indicated background and treatment. (**d**) Kaplan-Meier plots depicting the cumulative survival of the indicated mice. *p* value: statistical significance (Log-rank) for *Trp53*^null^ si*Wdr70* to the other two groups. (**e**) Examples of spontaneous tumours from terminally ill *Trp53*^null^ si*Wdr70* mice. (**f**) Immunofluorescence of Ki67 and HE staining for a representative region of liver hyperproliferation for the indicated genotypes. (**g**) HE (top) and IHC staining for Glypican-3 (bottom) in paraffin-embedded liver from the indicated genotypes biopsied at the indicated time. Nuclei were counterstained with haematoxylin. Magnifications are indicated on the left.

We next established cancer-prone animal models recapitulating the essential features of HBx-induced repair deficiency. *In vivo* RNAi of murine *Wdr70* (Extended Data Fig. 6c) was performed on *Trp53*^+/+^ and *Trp53* knockout (*Trp53*^null^) mice and these were compared to a mouse model with liver-specific expression of HBx under the control of *Albumin* promoter (_Albumin_*HBx*). Within two months of *HBx* expression or si*Wdr70* in *Trp53*^null^ mice, an increase in endogenous DSBs (53BP1 foci) and distorted nuclei were evident in liver tissue, especially in regions with severe pathological lesions (Fig. 6b,c and Extended Data Fig. 6d,e). When compared to *Trp53*^+/+^ cohorts, si*Wdr70* in *Trp53*^null^ dramatically shortened life span due to frequent malignancies in prolonged experiments (Fig. 6d,e).

We did not observe liver cancer within the timeframe of the si*Wdr70 Trp53*^null^ experiment, likely due to the prolonged timeframe required to develop HCC. However, liver biopsies from both si*Wdr70 Trp53*^null^ and _Albumin_*HBx* mice, when compared to *Trp53*^+/+^ si*Wdr70* or *Trp53*^null^ littermates, displayed hyperproliferation as revealed by Ki67 staining (Fig. 6f). The potential initiation of oncogenic processes that are likely to be prompted by the liver damage-repair cycle^48^ was evidenced by untimely expression of cytoplasmic Glypican-3 (Fig. 6g, lower panel), an oncofetal proteoglycan used as a biomarker for diagnosing early HCC^49^. We conclude that, by concurrent repressing CDW19S-ESM and p53, HBx is cancer-promoting.

### Targeting HBV-induced HRD suppresses the disease progression of HBVHCC

In the context of *BRCA1* ablation, the preferential channelling of DSB repair to error-prone NHEJ promotes the generation of toxic chromosomal structures that are exacerbated upon PARP inhibition^11,50^. We observed that chromosomal aberrations were synergistically exacerbated in *HBx*^EE^ and T43 cells upon PARPi (Olaparib) treatment and this was suppressed by si*53BP1* (Fig. 7a,b). Strikingly, HBV-positive cells (T43), when compared to HBV-free cells (L02), were sensitive to a range of PARPi (Fig. 7c and Extended Data Fig. 7a,b), indicating a strong SL pair of HBV/HBx and PARP inhibition. To establish if this was linked to CRL4^WDR70^, we knocked down *WDR70* in L02 and T43 cells (Fig. 7d). si*WDR70* sensitized L02 cells to PARPi, but showed no additional effect on T43 cell PARPi sensitivity, indicating CRL4^WDR70^ ablation by HBx contributes to HBV-dependent SL. Consistent with this, T43 cell PARPi sensitivity was rescued by sh*53BP1* (Fig. 7d). To extrapolate from cellular models, we established T43 xenografts in athymic nude mice. Tumour growth was strongly restricted by 131.5 mg/kg/day Olaparib mono-treatment (Fig. 7e and Extended Data Fig. 7c). Notably, Olaparib (41 mg/kg/day) and low doses of Cisplatin (0.42/mg/kg/2 days) acted synergistically on both cellular viability and T43 xenografts (Fig. 7f,g).

**Fig. 7.**
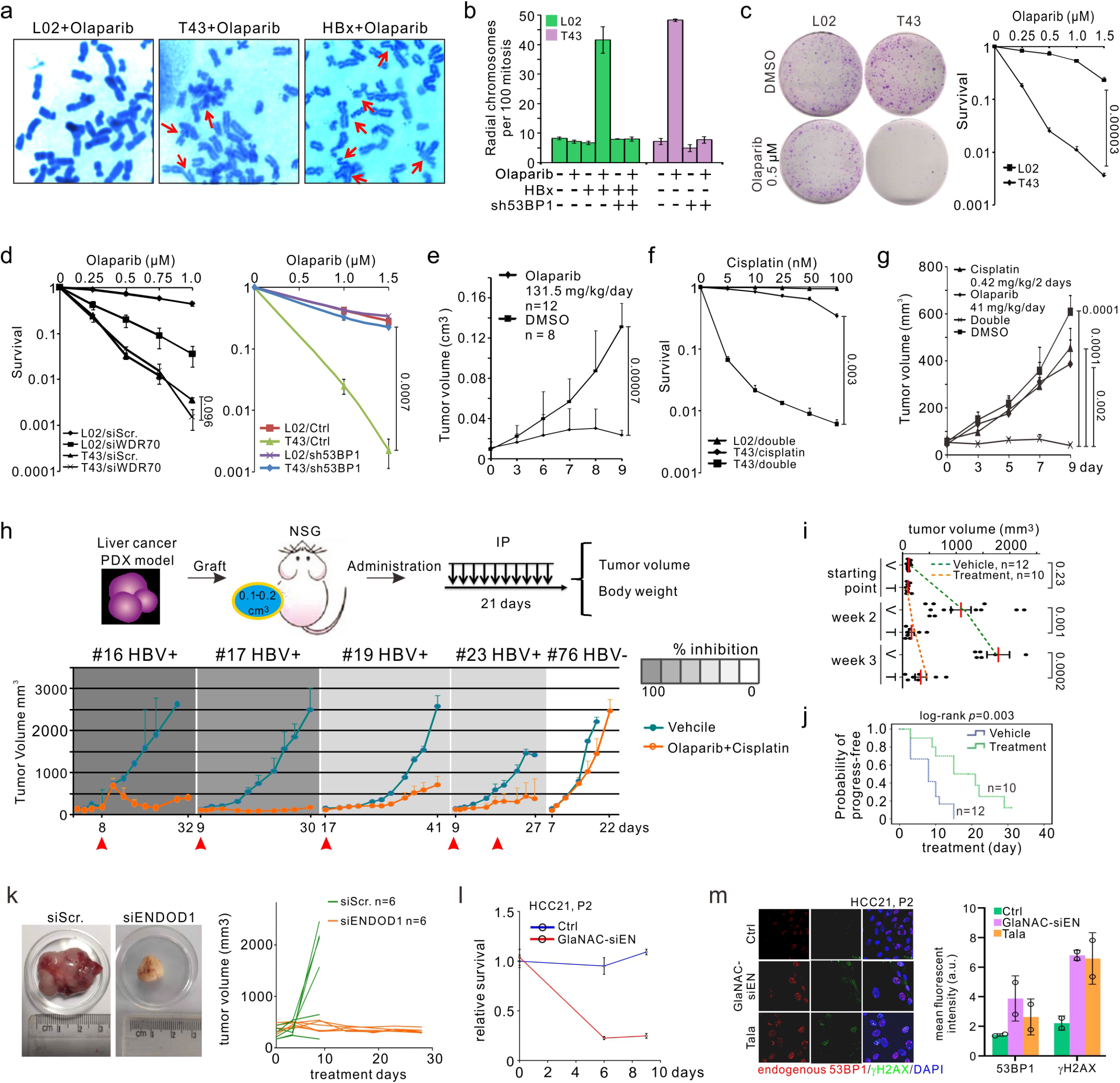
Curbing HBVHCC progression by HRD-targeting SL reagents. (**a**-**b**) Representative images (a) and quantification (b) of aberrant chromosomes in the indicated cells treated with Olaparib (1 μM) and/or sh*53BP1*. (**c**) Giemsa staining for colony formation (left) and survival curves (right) of control L02 (HBV-) and T43 (HBV+) cells mono-treated with Olaparib. Survival at end points was analysed for statistical significance. n = 3 biological repeats. Error bars = s.d. *t*-test. (**d**) Survival curves for L02 and T43 cells treated with si*WDR70* (left) or sh*53BP1* (right) with simultaneous exposure to indicated concentrations of Olaparib. (**e**) Responses of T43 xenografts to monotreatment with Olaparib or vehicle. Error bars: s.d, *t*-test. (**f**-**g**) Responses of T43 cells (f, 3 biological repeats) and xenografts (g, 6 littermates included) to conjunctive administration of Olaparib and Cisplatin. Tumour volumes were presented as means ± SD. DMSO: equivalent amount of solvent solution. Error bars: s.d, *t*-test. (**h**) Schematic of PARPi administration to HCC engraftment in NOD-SCID mice (top) and tumour responses (bottom). Tumour volumes of 4 HBVHCC (#16/17/19/23) that responded to therapy and 1 HBV-free HCC (#76: progressive disease) are shown at indicated days after inoculation. Olaparib: 33.3 mg/kg/day; Cisplatin: 0.5 mg/kg/2 days (O/C). Horizontal axis: days after tumor transplantation. Arrows: starting date of medication. (**i**) Tumor response for HBVHCC PDX sublines treated with vehicle or O/C at week 2-3. Graphs show mean ± s.e.m, analysed with two-sided unpaired Student’s *t*-test. (**j**) Kaplan-Meier plot indicating progression-free survival of HBVHCC sublines. The *y* axis is the percentage of animals whose tumor volumes were smaller than 300 mm^3^. *p* value was calculated by log-rank test. (**k**) Responses of T43 xenograft in naked mice to si*ENDOD1*. Representative tumours dissected at termination of experiments (left) and tumour volume (right) are shown. (**l**) Relative survival for primary HBVHCC culture (HCC21, passage 2) treated with GalNac-si*ENDOD1* or control siRNA. n = 3 biological repeats. Error bars = s.d. (**m**) Left: staining for endogenous 53BP1 and γH2AX in HCC21 treated with GalNAC-si*ENDOD1* or Talazoparib (20 nM, 48 hours). Right: quantification for fluorescent intensity. n = 2 biological repeats (lack of primary HCC cells), error bars: range.

The overall survival of HBVHCC patients is low and few contemporary chemotherapeutic treatments are widely applicable. To establish if PARPi could form a potential SL treatment for HBVHCC, patient-derived xenografts (PDX) were implanted in immunocompromised NOD*-Prkdc*^*scid*^-*IL2rg*^(em1 IDOM)^ mice and subsequently mice were treated with Olaparib at clinically relevant and mouse-equivalent doses (100 mg/kg/day)^51^ in conjunction with a low dosage of Cisplatin (0.5 mg/kg/2 days). Cisplatin-insensitive HCC xenografts from four HBV-positive patients (#16, 17, 19 and 23) and 1 HBV-free (#76) patient were subjected to one course of Olaparib/Cisplatin (O/C) conjunctive treatment. Body weights in both O/C and vehicle groups were monitored throughout the experiment (Extended Data Fig. 7d). Compared to the unrestricted tumour growth in the vehicle only group, all HBVHCC cases treated with O/C displayed tumour growth inhibition (TGI, from 95.17% to 51.09%) at terminal therapy (Fig. 7h), reflecting significantly delayed tumor progression (Fig. 7i and Extended Data Fig. 7e). Furthermore, O/C treatment produced a significantly longer median period of progression-free survival (PFS) compared to that of vehicle groups (*p*=0.003; Fig. 7j). In contrast, xenografts derived from HBV-free HCC (#76) exhibited no therapy response when subjected to the same course of O/C treatment (Fig. 7h).

We recently uncovered an atypical endonuclease (ENDOD1) that counteracts PARP trapping on damaged chromatin^26^ and demonstrated cell lethality upon concomitant ENDOD1-HR ablation. This potentially establishes a second therapeutic SL pair other than PARPi-HRD. We thus examined the SL between ENDOD1 and viral HRD. Monitoring cell proliferation and colony formation following siRNA of *ENDOD1* showed toxicity towards T43 but not L02 cells (Extended Data Fig. 7f,g). In T43 xenograft models, systemic *ENDOD1* knockdown yielded a progression-free outcome (Fig. 7k). N-acetylgalactosamine (GalNac)-modified siRNA is hepatocyte prone^52,53^ and *ENDOD1* ablation using this reagent prevented viability of primary HBVHCC cells (HCC21) and this correlated with increased γH2AX and 53BP1 foci formation at levels comparable to those seen following Talazoparib treatment (Fig. 7l,m). This suggests that inviability is attributable to DSB generation. Collectively, the response of HBVHCC to PARP or *ENDOD1* inhibition supports the therapeutic value of exploiting HRD to target this chemo-refractory cancer via SL.

## Discussion

In addition to regulating proteolysis, the 19S regulatory particle has a variety of non-proteolytic functions. Among these are a range of activities in the context of chromatin, including transcription initiation and elongation^18,54,55^, roles in histone modification^56^ and DNA repair^19-25^. In this study we identify a RP decorated with CRL4^WDR70^ (CDW19S) that assembles on chromatin in the vicinity of DSBs (see Fig. 4j). We show that CDW19S can be segregated into distinct functional modules defined by ubiquitin anchorage (PSMD4^Rpn10^) and subdomains required for the resection activity of MRE11 (MRM) and EXO1 (ESM) nucleases. Ablating subunits compromising the MRM resulted in resection defects at both 0.5 Kb (proximal) and 2.5 Kb (distal) locations from a DSB, whereas ablating ESM components exclusively caused distal resection defects. Interestingly, the ESM could be further subdivided into two discrete subunit groups influencing either distal POH1^Rpn11^ functions or CRL4^WDR70^ to coordinate the removal of the 53BP1 resection barrier protein.

Mechanistically, we identified ADRM1^Rpn13^ as a direct catalytic substrate for CRL4^WDR70^ associated with the RP and show that preventing ADRM1^Rpn13^ degradation by mutating a single lysine residue (K99>R) attenuates distal resection, thus identifying ADRM1^Rpn13^ as a key inhibitor of long-range resection that functions with 53BP1. Intriguingly, the equivalent K99 residue only exists in one of the fission yeast ADRM1 homologs, Rpn13b, and it was this protein that co-purified with WDR70 (cf. Extended Data Fig. 4g and Fig. 2a). This suggests a functional divergence between fission yeast Rpn13 homologs in the context of UPS and 19S RP chromatin functions in low eukaryotes, while in human cells all functions are carried out by a single ADRM1^Rpn13^ protein. We propose that CDW19S, upon association with chromatin that is ubiquitinylated by RNF8 at sites of DSB, engages its associated enzymatic activities (i.e. CRL4^WDR70^, POH1^Rpn11^) to stimulate nucleases and initiate resection in a BRCA1-dependent manner. Furthermore, we reconciled the functions of 19S in DNA repair and the regulation of damage-induced gene expression (exemplified by *p16*^INK4a^), by demonstrating that they both feature ubiquitin recognition via RNF8/PSMD4^Rpn10^ and CRL4^WDR70^ decoration of RP.

The HBV oncoprotein HBx plays multiple roles during HBVHCC development, including redirecting the Cullin4-DDB1 scaffold to promote the degradation of the SMC5/6 complex that counteracts viral replication^4^. By hijacking the DDB1 containing ubiquitin ligases, HBx causes a deficit in the cellular ligases that rely on the Cullin4-DDB1 scaffold, including CRL4^WDR70^(ref^5^). The ablation of CRL4^WDR70^ results in a defect in homologous recombination^5^ and renders HBV infected cells homologous recombination defective (HRD). Here we have identified the underlying mechanism by which HBV/HBx pathogenesis causes HRD and carcinogenesis. By compromising the integrity of the CRL4^WDR70^, we demonstrate that HBx prevents the integration of Cullin4-DDB1 into chromatin-associated CDW19S. In the absence of its Cullin scaffold, WDR70 associated with this torso-CDW19S is unable to promote ADRM1^Rpn13^ degradation and thus cannot relive the barrier to EXO1-dependent resection, thus compromising HR efficiency.

Using mouse models we demonstrated that CRL4^WDR70^ ablation, when combined with compromised p53, leads to carcinogenesis. This confirms that the combined action of HBx in forming torso-CDW19S and preventing p53 transcriptional transactivation^57^ is capable of driving cancer development in a manner resembling that previously characterised for *BRCA1*-mutated cancers. This underlines the importance of dual inhibition of CDW19S-ESM and p53 in driving HBVHCC carcinogenesis. Further elucidating distinct biochemical activities by which CDW19S regulates the choice between NHEJ and HR and how this is regulated by BRCA1 will be critical to fully understand this viral HRD subtype and thus precisely how HBVHCC occurs.

PARPi has been successful in treating breast cancers manifesting with BRCAness HRD^58-60^. We showed that HBV (HBx)-induced HRD is similarly synthetic lethal to PARP inhibition. We further demonstrate, using Xenograft models of T43 cells or patient-derived HBVHCC materials, that treatment with PARPi alone, or treatment with a combination of low dose cisplatin and PARPi significantly constrained tumour growth. This underlines the potential for SL treatment using PARPi. In previous work we identified the atypical nuclease ENDOD1 as a protein required in human cells to reduce the levels of trapped PARP and showed that *ENDOD1* ablated cells were synthetic lethal with homologous recombination defects^26^. Here we show knockdown of *ENDOD1* causes increased levels of DNA damage and forces cytotoxicity in the context of *HBx* expression and, using xenografts, show that knockdown of *ENDOD1* by whole animal siRNA results in reduced tumour growth. This indicates that, in addition to PARPi, future reagents targeting ENDOD1 have potential for advancing therapy against HBVHCC.

Taken together, we identify the mechanism by which HBx causes homologous recombination defects, demonstrate that the combination of HRD and p53 deficiency resulting from HBx expression can account for the oncogenic nature of HBV infection and establish that HBVHCC is an HRD cancer. Our findings provide a mechanistic and pre-clinical justification for targeting HBVHCC by SL.

## Acknowledgements

This work is supported by the National Natural Science Foundation of China (31771580, 82073027, 81702795, 81972592, 81821002), Sichuan University (2020SCUNL204, SCU2019C4198), Department of Science and Technology of Sichuan Province (2019JDTD0020, 2020YFH0019), and the 1.3.5 project for disciplines of excellence of West China Hospital (ZYGD20007, ZYJC18011, ZYGD18003 and ZYJC21021). A.M.C. acknowledges MRC award G1100074. There are no conflicts of interest in this study.

## Author contributions

C.L. purified CDW19S complex and M.Z. characterised it. L.R. and H.W. analysed 53BP1 and BRCA1 in the absence of CRL4^WDR70^. J.C. assisted I-*Sce*I repair assay. X.W. performed immunoblotting and cell culture. W.Z., Z.L. and J.J. identified components by mass spectrometry. X.M. advised on *TP53* experiments. Z.T. contributed to anti-cancer studies. X.M. and J.H. contributed ADRM1 degradation. D.K. A.M.C. and C.L. directed and interpreted DDR analysis. C.L. and A.M.C. wrote the manuscript.

