## Supplementary data for "Mechanistic basis for targeting homologous recombination defective liver cancer via synthetic lethality"

**Extended Data Fig. 1 HBx disturbs the CRL4$^{WDR70}$-dependent 53BP1-BRCA1 balance at IRIF.**
(**a-b**) Quantification of 53BP1 IRIF over time in L02 cells treated with *DDB1* or *WDR70* siRNA (a), or *HBx*-expressing lentivirus (b). (**c**) Quantification of 53BP1 foci with an obvious cavity at 6 hours after IR in the indicated cells (pLV and HBx: lentivirus vector control or expressing HBx in L02 cells). (**d**) Representative image (left) and quantification (right) of co-localised RPA32 (green) and 53BP1 (red) IRIF at indicated time post irradiation in lentivirus vector control (pLV) or lentivirus expressing HBx in L02 cells. Note that the percentage of double positive foci decreased with time in control L02 cells but increased over time in the presence of HBx. Cells were pre-extracted with 0.1% Triton- X100 to visualize RPA32. (**e**) Quantification for pRPA32 and RAD51 IRIF in 293T cells treated with si*WDR70* or *HBx* expression. *53BP1* was knocked down by infecting with sh*53BP1* lentivirus. pLV: vector control. (**f**) Enumeration of pBRCA1 IRIF in the indicated cells or L02 cells with the indicated treatments (pLV and HBx: lentivirus vector control or expressing HBx in L02 cells). All experiments in the figure: n = 3 biological repeats. Error bars = s.d. *p* values by students *t*-test.

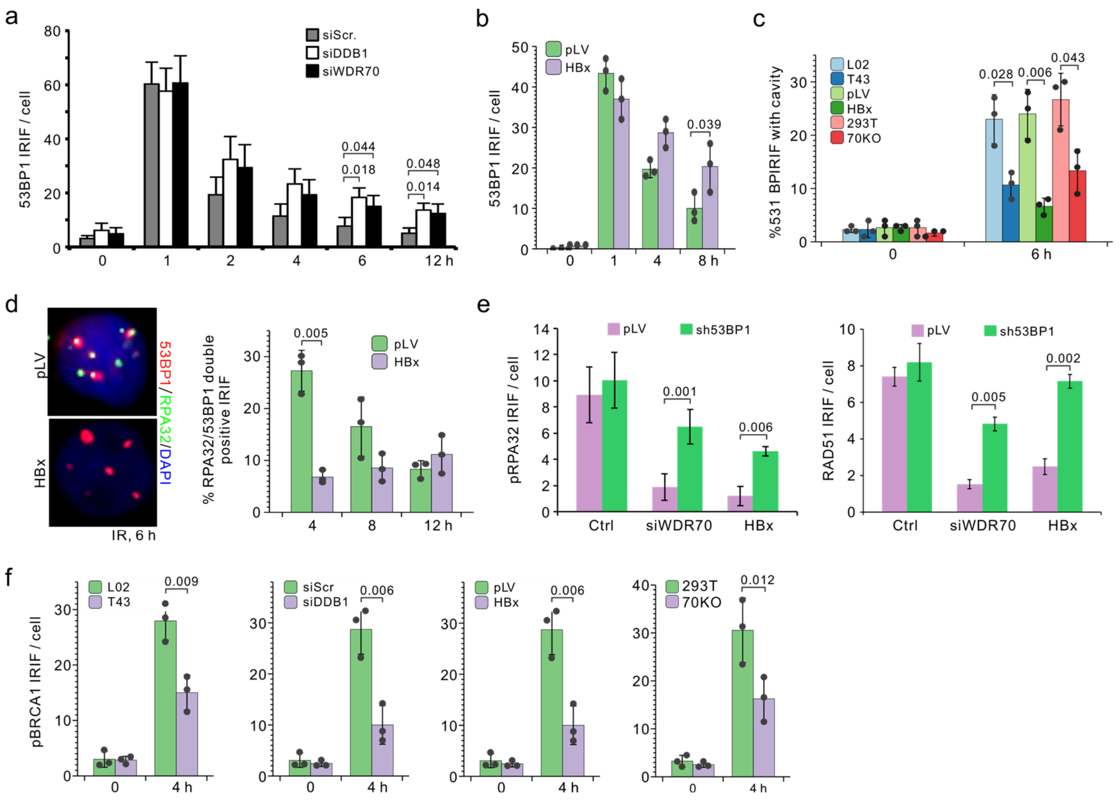

**Extended Data Fig. 2 19S RP recruits CRL4$^{WDR70}$ to DSBs.**

(**a**) co-IP of endogenous WDR70 and DDB1 by Flag-tagged 19S subunits. (**b**) co-IP of endogenous WDR70 by Flag-tagged PSMD2$^{Rpn1}$, PSMD4$^{Rpn10}$, POH1$^{Rpn11}$ and PSMC1$^{Rpt2}$ in chromatin extraction pre-treated with CPT (2 μM, 2 hours) or not. (**c**) co-IP of endogenous WDR70 and DDB1 as above with Flag- PSMD4$^{Rpn10}$. Pre-treatment with si*WDR70* or si*DDB1* as indicated. (**d**) Immunoblotting for pRPA32 and H2B monoubiquitination (uH2B) in L02 cells with indicated siRNA and CPT treatment. (**e**) *Asi*SI-dependent DSB-association of pRPA32 (chromosome 1: 89,458,595 - 89,458,603) assayed by ChIP 4 hours after 4-OHT addition. *WDR70* was depleted by siRNA. n=3 biological repeats. Error bars; s.d. (**f**) Left: illustration for *Xba*1-based resection assay at the selected *Asi*SI-dependent DSB site. Right: example of monitoring ssDNA production by semiquantitative PCR. (**g**) Knockdown efficiencies in DIvA cells evaluated by RT-qPCR for individual 19S genes relative to si*Scramble* after normalization to 18S *rRNA* levels. Semi-quantitative PCR for *PSMC5*$^{Rpt6}$, *PSMC6*$^{Rpt4}$ and *PSMD12*$^{Rpn5}$ as well as actin controls are shown in inset. n=3 biological repeats. Error bars; s.d. (**h-i**) ChIP assay for Flag-tagged 19S subunits (h) or PSMD4$^{Rpn10}$ (i) at indicated distances from an *Asi*SI-dependent DSB with indicated treatments.

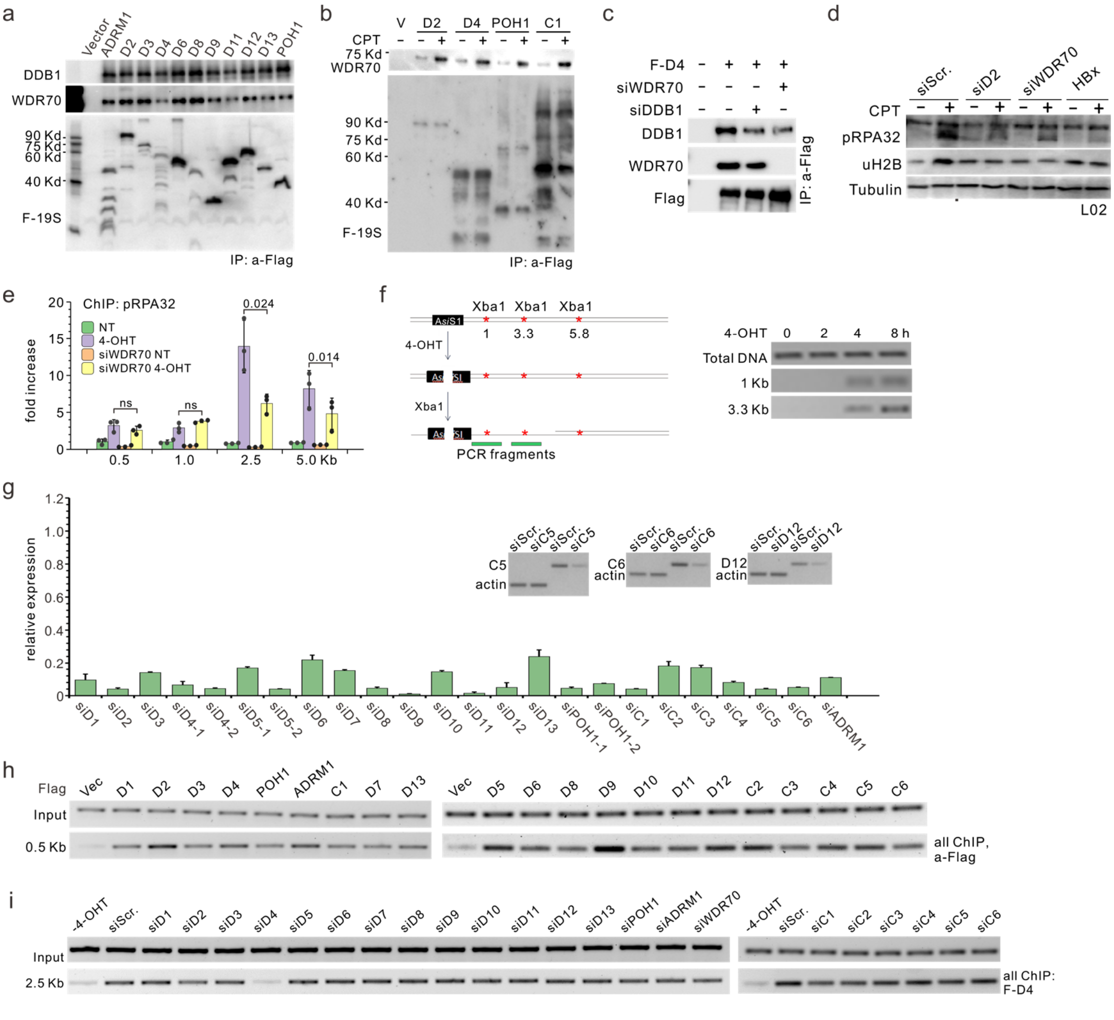

**Extended Data Fig. 3 Modular functions of CDW19S.**

(**a**) DIvA cells transfected with two different siRNA each for *PSMD4*$^{Rpn10}$ and *PSMD5*$^{Hsm3}$. The DSB loading of pRPA32 was quantified at 2.5 Kb from an *Asi*SI site in the presence or absence of concomitant expression of siRNA resistant plasmids. (**b**) pRPA32 loading at 0.5 Kb from an *Asi*SI-dependednt DSB upon *PSMD2*$^{Rpn1}$, *PSMD4*$^{Rpn10}$, *PSMD7*$^{Rpn8}$ and *POH1*$^{Rpn11}$ depletion in the presence or absence of concomitant si53BP1. (**c,d**) Quantification (c) and representative images (d) for MRE11 IRIF at the indicated time points after siRNA treatment for selected CDW19S genes. (**e-f**) Coomassie staining showing purified recombinant WDR70 (0.5 µg) from insect cells and PSMD5$^{Hsm3}$ (0.5 µg) from *E. coli* with indicated tags (e), plus purified commercial 19S proteasome (1 µg, R&D, E-367, f). (**g**) Left: Immunoblotting showing the antibody specificity of $\alpha$-PSMD5$^{Hsm3}$ in 293T cells with pre-treatment, or not, of si*PSMD5*$^{Hsm3}$. Right: immunoreaction detected by $\alpha$-PSMD5 in purified19S particles. (**h**) ChIP for Flag-KU80 at 0.5 Kb from an *Asi*SI-dependent DSB following the indicated siRNA treatment. (**i**) IF for 53BP1 30 min or 8 h after ionizing irradiation with silencing, or not, of POH1$^{Rpn11}$. Representative images (left) and quantification for fluorescent intensity (right) were shown. For all graphs: n = 3 biological repeats. Error bars = s.d. *p* values by *t*-test shown for indicated groups. ns: no significant difference.

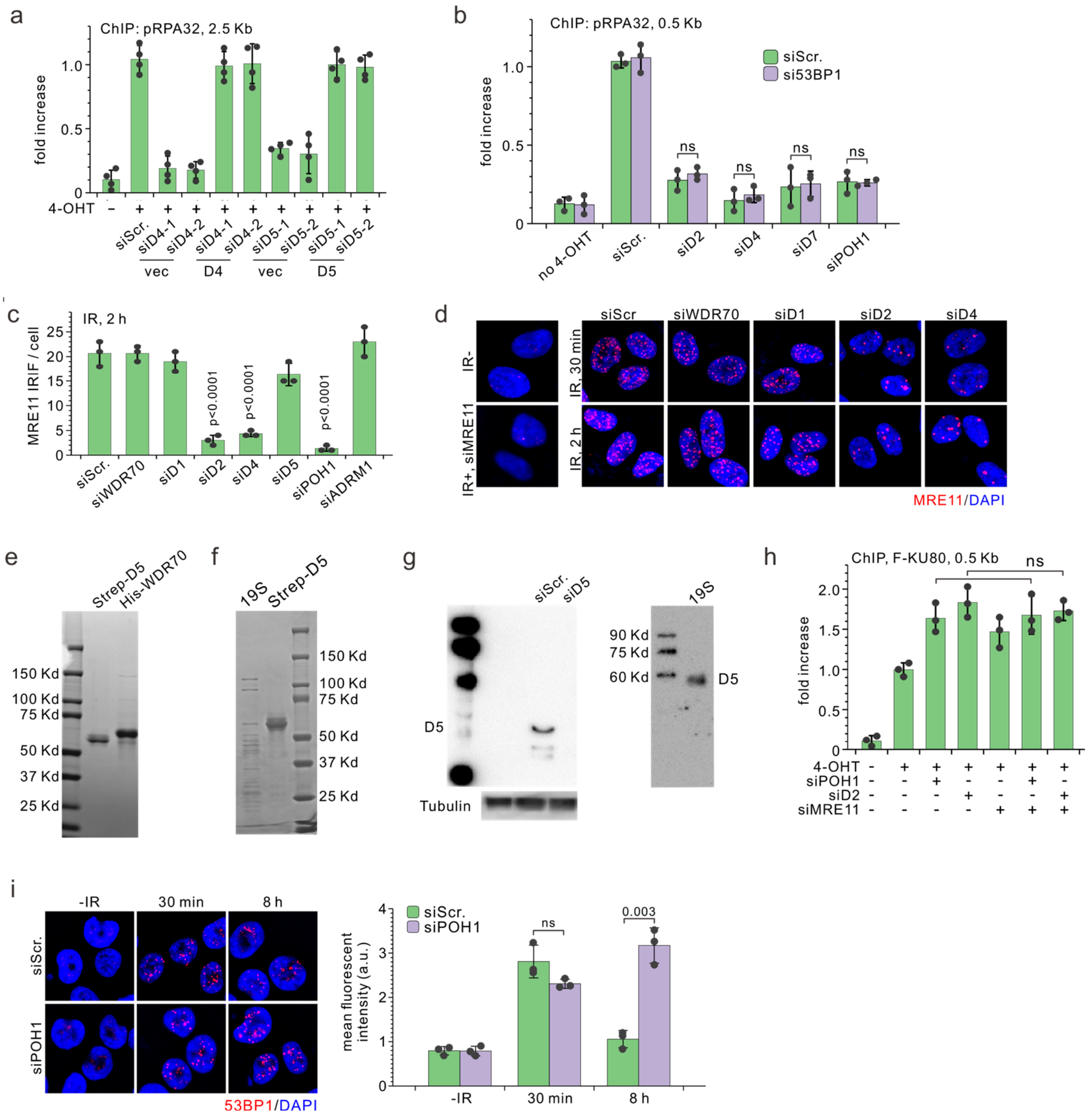

**Extended Data Fig. 4 ADRM1$^{Rpn13}$ as ubiquitin target of CRL4$^{WDR70}$.**

(**a-b**) ChIP for ADRM1$^{Rpn13}$ (a) and EXO1 (b) at the indicated distances from an *Asi*SI-dependent DSB in the absence of *WDR70*, *ADRM1*$^{Rpn13}$, or both. (**c**) Schematic for ADRM1$^{Rpn13}$ protein functional domains (left) and ubiquitin binding assay (right) for ADRM1$^{Rpn13}$ with biotinylated ubiquitin monomer, K48 and K63 chains (0.5 µg of each) immobilized on streptavidin beads. Flag-tagged ADRM1$^{Rpn13}$ (wild type and mIFD) was purified from 293T cells and monitored by immunoblotting with $\alpha$-Flag. (**d**) Chromatin fraction from CPT-treated cells expressing full length (FL) Flag-ADRM1$^{Rpn13}$, or mutants with deleted Pru or DEUBAD domains. T: total extracts; C: chromatin. (**e**) Left: ChIP assay at 2.5 Kb from an *Asi*SI-dependent DSB for Flag-tagged ADRM1$^{Rpn13}$. Right: expression of wild type and mIFD monitored by immunoblotting with $\alpha$-Flag. (**f**) Left: representative immunoblotting measuring levels of ADRM1$^{Rpn13}$ following CPT treatment as for Fig. 4c. Right: immunoblotting for ADRM1$^{Rpn13}$ in the presence of MG132 and/or CPT. Cells were pre-treated, or not, with si*WDR70*. (**g**) Alignment of human and *S. pombe* ADRM1$^{Rpn13}$ homologues, positions of conserved lysines are labelled. Note that only the human and yeast Rpn13b contain the K99 residue. (**h**) Ub pulldown assay in indicated cells for ubiquitinated Flag-ADRM1$^{Rpn13}$ upon expression ADRM1 K99-only protein. (**i**) ChIP assay at 2.5 Kb from an *Asi*SI-dependent DSB for wild-type, K99R and K99-only Flag-ADRM1$^{Rpn13}$ in DIvA cells treated, or not, with si*WDR70*. (**j-k**) Coomassie staining for purified CRL4$^{WDR70}$ tetramer from sf9 cells (j) and Flag- or GST-Flag-ADRM1$^{Rpn13}$ from *E. coli* (k). (**l**) Reconstitution using indicated E2 enzymes. Ubiquitinated ADRM1$^{Rpn13}$ was monitored by streptavidin for biotinated ubiquitin. For all graphs: n = 3 biological repeats. Error bars = s.d. *p* values by *t*-test shown for indicated groups. ns: no significant difference.

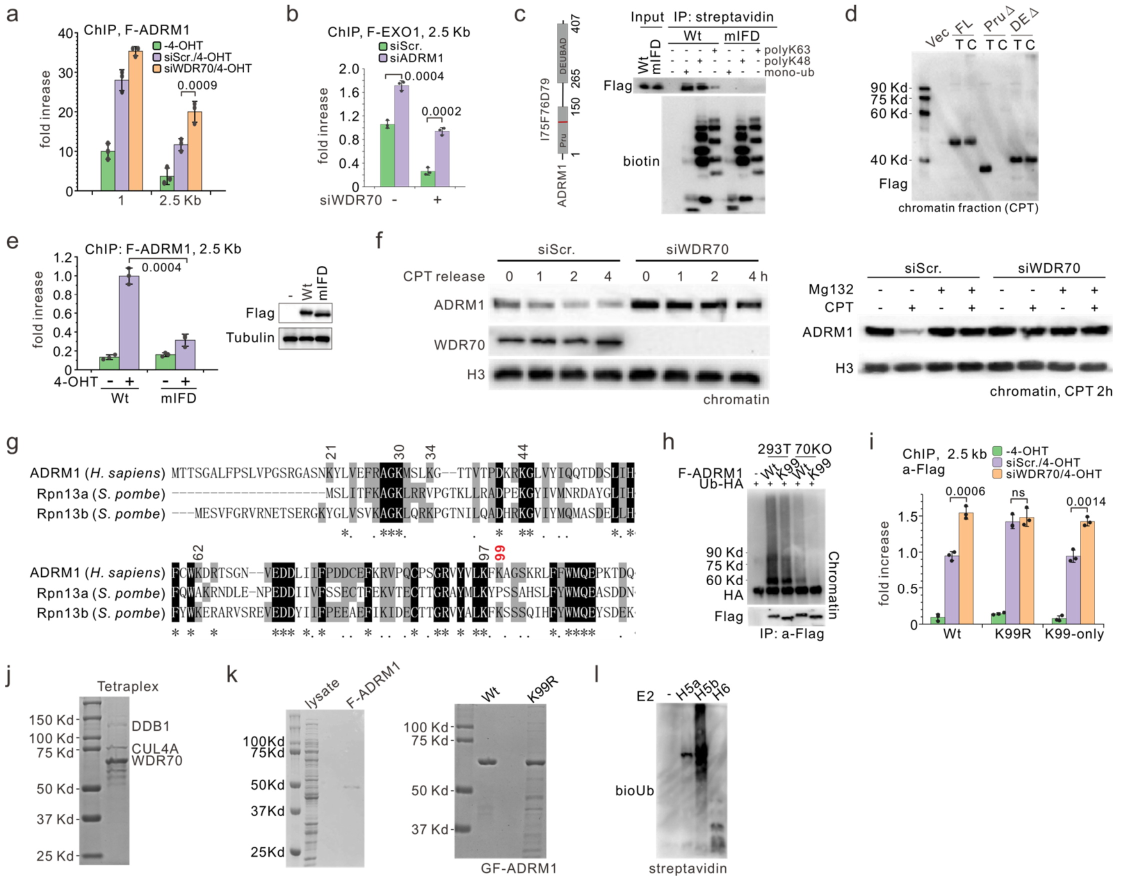

**Extended Data Fig. 5 HBX interferes with the organization of CDW19S.**
(**a**) co-IP of endogenous DDB1 by Flag-tagged WDR70 in L02 and T43 cells. si*HBx* was introduced to restore interaction in T43 cells. (**b-c**) ChIP assays using CRISPR-induced DSB showing the dependence of WDR70 loading to DSB on 19S RP but not vice versa (as for PSMD4$^{Rpn10}$). These results reproduce the observation in the DIvA system. (**d**) Quantitative PCR (top) and immunoblotting (bottom) showing the expression of indicated repair factors in DIvA cells upon depleting *WDR70*, *PSMD4*$^{Rpn10}$ or *PSMD5*$^{Hsm3}$. Two primer sets (a,b) were used for quantifying the *BRCA1* mRNA. (**e-g**) *p16*$^{INK4a}$ expression evaluated by immunoblotting (e) or RT-qPCR (f,g) in HBV-bearing (HepaG2.2.15 or T43), *HBx*$^{EE}$ or *WDR70* ablated L02 cells upon CPT treatment (0.4 μM, 12 hours). *p16*$^{INK4a}$ mRNA levels were normalized with internal control (GAPDH) and displayed relative to the si*Scramble* with CPT treatment. (**h-i**) As above, CPT-induced *p16*$^{INK4a}$ mRNA in L02 cells was assessed following the indicated treatments. Knockdown efficacy of *RNF8* is shown on the right panel of (i). (**j-k**) ChIP for DDB1/WDR70 (j) and H2B monoubiquitination (k) at α or β sites of the *p16*$^{INK4a}$ promoter in L02 cells subjected to the indicated treatments. For all graphs: *p* values by *t*-test shown for indicated groups. n = 3 biological repeats. Error bars = s.d.

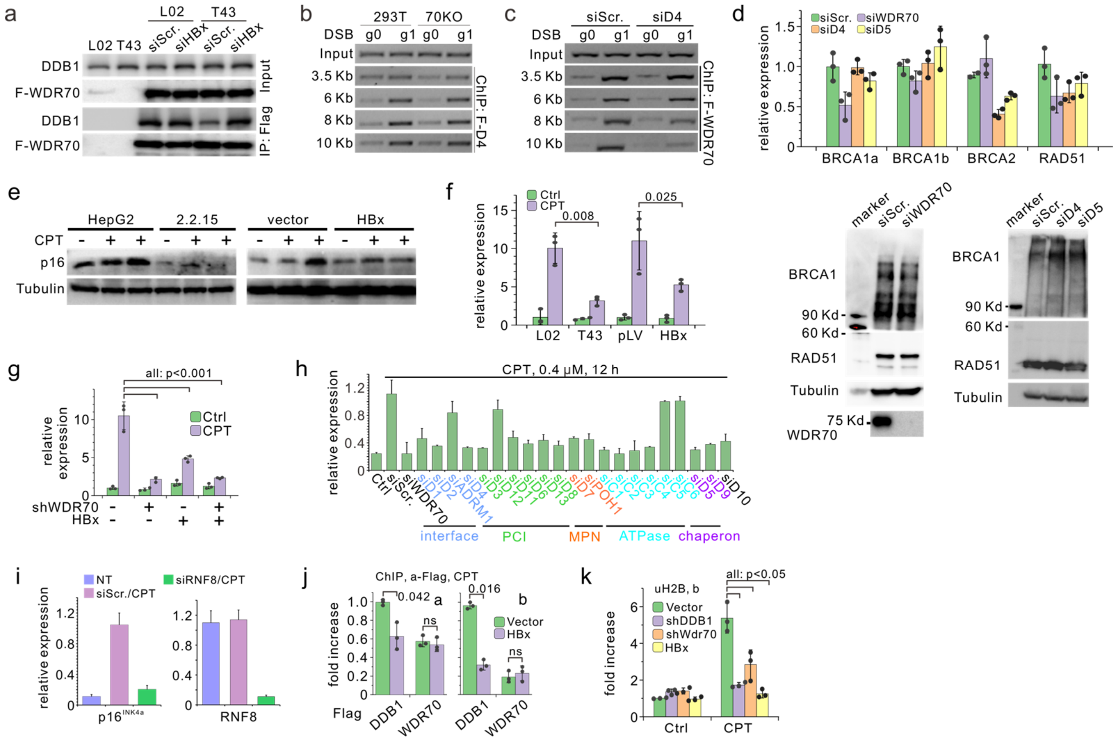

**Extended Data Fig. 6 Simultaneous suppression of *WDR70* and *TP53* by HBx.** (**a**) Semi-quantitative PCR for death targets of p53 in the presence of HBx in HBV-free cells (L02 and HepaG2) or in HBV carrier cells (T43 and HepG2.2.15). si*WDR70* did not impact on the p53 targets at the level of transcription. (**b**) Colony formation assays showing sensitivity to cisplatin only or double treatment with cisplatin and 50 nM of the p53 activator Nutlin (Double). n = 3 biological repeats. *p* values by *t*-test shown between single and double treatment groups of T43. Error bars = s.d. ns: no significant difference. (**c**) Verification of the silencing of murine *Wdr70* in liver by semi-quantitative RT-PCR a week after *in vivo* RNA injection (2.5 mg/Kg). si*Wdr70*-001 was used for following animal experiments (2.5 mg/Kg/injection, twice a week). (**d-e**) Quantifications for 53BP1 foci (d) and nuclear morphology (e) as in Fig. 6b and 6c. 53BP1 foci were quantified either throughout the liver (pan-liver) or in regions displaying severe pathological changes (patho. lesion). *p* values by *t*-test are shown for indicated groups. n = 3-4 biological repeats from individual mouse tissues. At least 50 cells were counted for each repeat. Error bars = s.d.

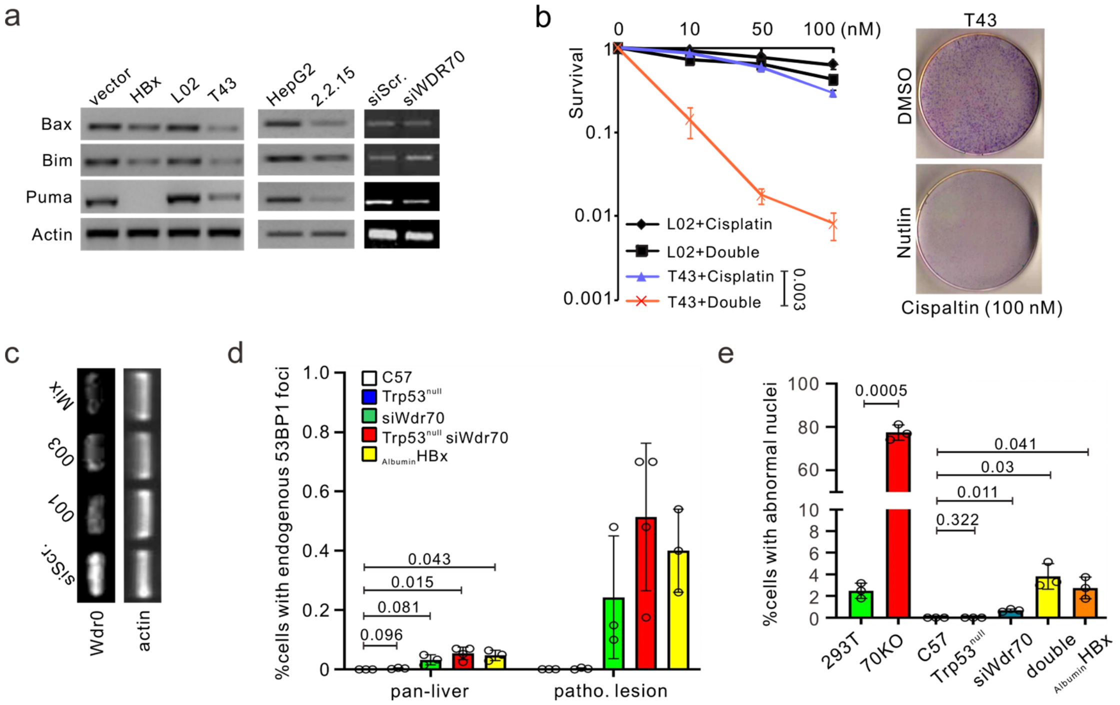

**Extended Data Fig. 7 Targeting HRD curbs the disease progression of HBV-positive tumors.**

(**a**) Viability of HepG2 (HBV-) and HepG2.2.15 (HBV+) upon Olaparib treatment. n = 3 biological repeats; *p* values by *t*-test are shown; error bars: s.d. (**b**) Percentages of T43 survival after challenging with indicated PARPi assayed by colony formation. n = 3 biological repeats; *p* values by *t*-test are shown; error bars: s.d. (**c**) Dissected tumour (left) and average weight (right) of T43 xenografts obtained at endpoints of experiments in Fig. 7e. (**d-e**) Same experiments as in Fig. 7h showing longitudinal individual-body-weight curves from tumour-burdened mice (d), or individual tumour-growth curves (e) of HBVHCC xenograft sublines in vehicle (green) or Olaparib + CTP (OC: orange) treated groups. (**f**) Colony formation assay for L02 and T43 cells with indicated si*ENDOD1* or Olaparib (0.5 μM) treatments for 16 days. (**g**) Cell proliferation was quantified by haemocytometer cell counting for indicated time points. L02 or T43 cells were treated with control siRNA or si*ENDOD1* every 3 days. n = 3 biological repeats; error bars: s.d.

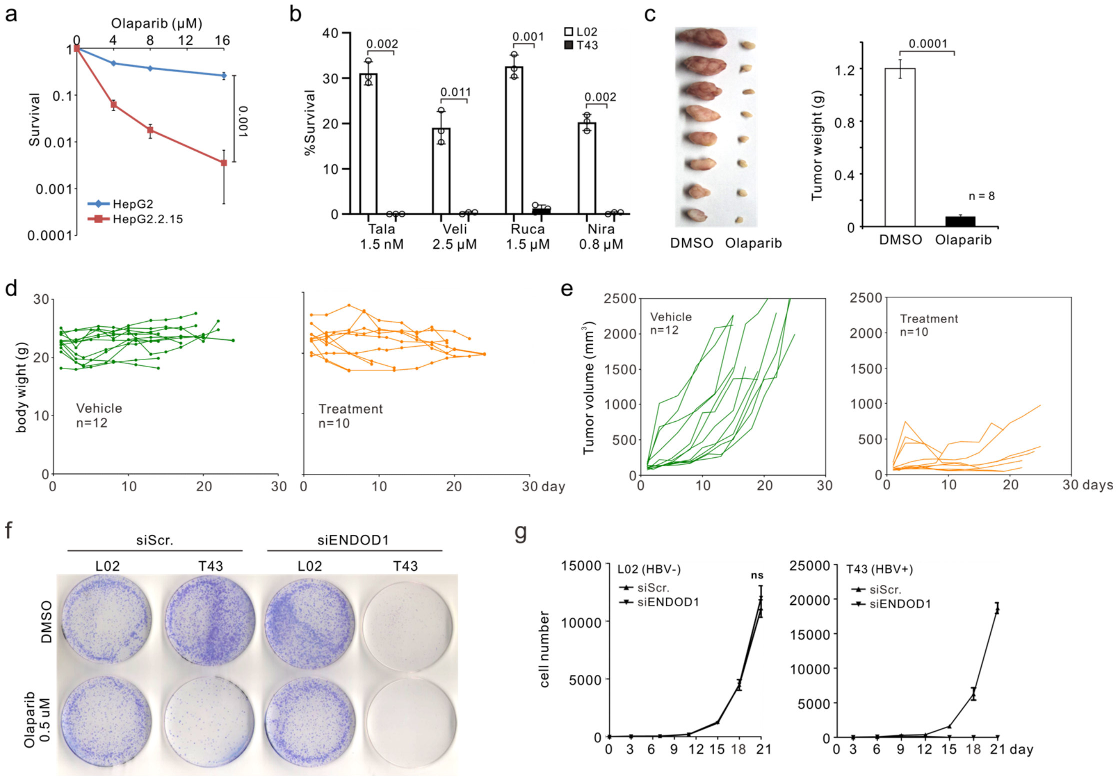

### Methods

### Materials and Methods

#### Cell culture and reagents

Human cell lines (HEK293T, *WDR70*$^{KO}$, RPE1, DIvA, HepG2 and HepG2.2.15, L02 and T43) were maintained in culture media supplemented with 10% foetal bovine serum. T43 cells with stably integrated HBV genomes were regularly selected in G418 (200 μg/ml) and examined for titration of HBs and HBe antigens by ELISA[1]. Cell lines were purchased from National Infrastructure of Cell Line Resource (Shanghai). All cell lines tested negative for mycoplasma contamination. Plasmid and siRNA transfections were carried out using Lipofectamine 3000 (Invitrogen, L3000015). Primary liver cancer cells were isolated from fresh cancer tissue using a cell scraper and cellular debris was removed by centrifugation. Cell pellets were resuspended in culture medium (Procell, CM-H140) and growth status was observed after 7-14 days. Primary cells were passaged and experiments were performed when cells grew ~80% confluent of plates. DSBs in DIvA system were induced by 300 nM 4-OHT (Sigma, H7904) for 4 h. For ATM kinase or proteasome inhibition, 10 μM KU55933 (Selleck, S1092) or 100μM MG132(Selleck, S2619) was used for pretreatment of 2 h.

#### Plasmids

For cloning of lentiviral vectors for expression of Flag-tagged proteins, PCR fragments were inserted into the *Eco*R I site of pLVX-G-Flag plasmid, using the In-Fusion cloning kit (Clontech, 639650). Plasmids used in this study are listed in Table S2.

#### Evaluation of gene expression

For detection of gene expression, $10^6$ cells were harvested and total RNA was extracted using NucleoZOL Reagent (740404, MNG). cDNA was obtained from 1 μg RNA by Reverse Transcription System (Promega, A3500). Real time-qPCR and semi-quantitative PCR were performed on Bio-Rad CFX96 Real Time System according to instruction of qPCR Kits (Promega, A6001) with three technical replicates for each sample. For RT-qPCR, the relative expression was calculated by fold change of target gene normalized to GAPDH/18S rRNA in each sample and the experimental group was normalized to the average of the control. PCR products for semi-quantitative were resolved by 2% agarose electrophoresis.

#### TAP-affinity purification for Wdr70 associating proteins

To purify proteins in complex with Wdr70 of *S. pombe*, 8 liters of *spWdr70* TAP-tagged or control strains were grown in YEP rich media. The cell pellet was washed and resuspended in an equal volume of lysis buffer (25 mM Tris-HCl, pH 7.5; 15 mM

EGTA, pH 7.5; 15 mM $MgCl_2$; 0.1% NP-40; 1 mM DTT; 0.1 mM NaF) supplemented with protease inhibitor (Complete$^{TM}$ Mini, Roche) plus DTT (1μM) and subjected to mechanical beating with glass beads in a Ribolyzer. The granular cell pellets and the powder of ground cells were be frozen at –80 °C and stored (up to 3 months) before use. Soluble extracts were clarified by centrifugation before binding to rabbit IgG-coated Dynabeads® M280 (Invitrogen), followed by extensive washing and cleavage by TEV protease. Purified Wdr70-associating components were resolved by 8-20% gradient SDS-PAGE and stained with Brilliant Blue G-Colloidal Concentrate Kit (Sigma, B-2025).

Excised gel bands were washed with 25 mM $NH_4HCO_3$ containing 50% Acetonitrile and dehydrated with 100% Acetonitrile, followed by treatment with 10 mM DTT and incubation for 1 h at 65°C. After cooling, gel samples were alkylated with 55 mM Iodoacetamide for 45 min at room temperature. Samples were digested with trypsin (1:50, w/w) dissolved in 25 mM $NH_4HCO_3$ at 37°C overnight. Digested peptides were extracted with 5 mM octyl-β-D-glucopyranoside in 0.25% Trifluoroacetic acid for 60 min at 37°C and directly applied onto the AnchorChipTM target (Bruker Daltonics) that was loaded with α-cyano-4-hydroxycinnamic acid (CHCA) thin layer matrix. Mass spectra of extracted peptides for each sample were determined using Ultraflex MALDI-TOF/TOF mass spectrometry (Bruker Daltonics) in a positive ion reflector mode. The ion acceleration voltage was 25 kV. Both MALDI-TOF spectra and the MS/MS spectra were processed by FlexAnalysis 2.2 and Biotool™ 2.2 and automatically searched against the Swiss-Prot database using Mascot software. Main parameters were set as follows: mass range from 800 to 4000 Da; S/N ≥ 3.0; fixed modification, carbamidomethyl (Cys); variable modification, oxidation (Met); maximum number of missing cleavages, 1; MS tolerance, 50 ppm; and MS/MS tolerance, 0.7 Da.

**Protein purification and pull-down assay**

Purified 6xHis-WDR70 (112-654 aa) and CRL4-WDR70 tetraplex from Sf9 insect cells were purchased from HitGen. For PSMD5 purification, the coding sequence was amplified from a cDNA library and cloned into the pET28a-2xStrep plasmid. For bacterial expression, BL21 strain transformed with expression vector was induced with 1 mM IPTG overnight at 16°C. 500 mL cells were pelleted and resuspended in 20 mL TBST buffer with protease inhibitor cocktail and sonicated. Centrifuged supernatant was incubated with MagStrep XT Beads (2-4090-002) and eluted with BXT buffer (100 mM Tris[pH=8.0], 150 mM NaCl, 1 mM EDTA, 50 mM biotin) for 1 h. For ADRM1 Wt or K99R purification, pET28a-Flag plasmids were used with otherwise similar protocol except incubation was with GSH beads (BeaverBeads, 70601-5) and elution with Buffer B (50 mM Tris-HCl, 10 mM glutathione).

For immunoprecipitation, $10^6$ cells were harvested and lysed in 200 μl of buffer A (10 mM HEPES [pH 7.9], 10 mM KCl, 1.5 mM $MgCl_2$, 0.34 M sucrose, 10% glycerol, 1 mM dithiothreitol, 0.1% Triton X-100 and protease inhibitors). Nuclei were collected in the pellet by low-speed centrifugation (1500 × g, 4 min, 4 °C) and further incubated with buffer B (2 mM EDTA, 0.2 mM EGTA, 1 mM dithiothreitol, and protease inhibitor mixture) for 10 mins on ice. After centrifugation (2000 ×g, 4 min), the pellet was resuspended in IP2 buffer (50 mM Tris [pH 8.0], 150mM NaCl, 1% NP40, and protease inhibitor cocktail) with100U Micrococcal Nuclease/DNase I and incubated on ice for 30 min to digest genome DNA. The chromatin fraction was clarified by high-speed centrifugation (21000 ×g, 10 min) and then incubation with pre-washed Flag M2 beads (Sigma, M8823) to pull-down Flag-tagged immunocomplex. Proteins were detected by Immunoblotting.

**Immunostaining**

Indirect immunofluorescent staining was described elsewhere [2]. Briefly, cryosectioned tissues or cells on coverslips were fixed with 4% PFA. Permeabilized sections were incubated with primary antibodies and labelled with secondary antibodies (anti-rabbit-Cy3 or anti-mouse-FITC), followed by mounting with anti-fade medium containing DAPI and visualising using Leica fluorescence microscope (DM4 M) or Olympus fluorescence microscope (BX51). For immunohistochemical staining (IHC), paraffinized tissues were deparaffinized and treated with citrate-based solution for antigen unmasking, and with $H_2O_2$/methanol solution for quenching endogenous peroxidase. Slides were incubated with primary antibody and visualised by ABC-DAB staining (Vector laboratory). Nuclei were counterstained with hematoxylin. All quantitative immunostaining analysis was performed by counting 100-200 cells from 3 independent experiments.

For measurement of 53BP1 exclusion from the core IRIF, images were processed using FV10-ASW 3.1 Viewer software, and the cavity of 53BP1 and sizes of pRPA32 foci was gauged by measuring pixel densities by ImagePro Plus 6.0 across the centre lines of foci with best visualisation. 50 - 200 foci were measured for each group.

**In vitro ubiquitination assay**

Recombinant E2 enzymes (R&D, K-980B) and ubiquitin conjugation initiation Kit (R&D, K-995) were used for ubiquitination assay. In brief, 500 ng purified ADRM1 or mutant protein was incubated with UBE1, UbcH5b, ubiquitin, $CRL4^{WDR70}$ tetramer

(500 ng) at 37°C for 1 h. The samples were mixed with SDS sample buffer and detected by Immunoblotting following SDS-PAGE. Recombinant ubiquitin mutants K48R (R&D, UM-K48R) and K63R (R&D, UM-K63R) were used for identification of ubiquitin chain species. Recombinant 19S proteasome with Flag-tagged UCH-L5 was purchased from R&D system (E-367).

**Measurement for efficacies of DSB repair**

For plasmid-based DSB repair system3, pCMV plasmids containing the I-*Sce*I restriction site were subject to thorough in vitro digestion. Cut plasmids (NHEJ and SSA: 5 μg, HR: 10 μg) were transfected into 293T or WDR70 knockout cells and allowed to repair in vivo for 48 hours. Recombined or ligated plasmids by SSA/HR (primers 3/4 or NHEJ (primers 5/4) were recovered from cells by HighPure PCR Template Preparation Kit (Roche, 11796828001), followed by quantitative Real-time PCR with appropriate primers (Table S3) specific for SSA/HR or NHEJ fragments using a Bio-Rad CFX96 Real Time System. The amount of extracted plasmid DNA was normalized to the product of primers 1/2. The relative frequency of each repair pathway was defined as fold changes relative to qPCR values of repaired DNA extracted in parallel from wild type or mock infected 293T cells.

For the *Xba*I-dependent resection assay, genomic DNA from 2 x $10^5$ DIvA cells following 4-OHT induction was purified using High Pure PCR Template Preparation (Roche, 11796828001). For each sample, 300 ng of extracted DNA was digested with *Xba*I at 37 °C for 4 h and the reaction was stopped by heating at 65 °C for 10 mins. For ssDNA quantification, 20 ng digested sample was amplified by qPCR using primers listed in Extended Data Fig. 2f and Table S3.

**Chromatin immunoprecipitation (ChIP)**

ChIP assays were performed as previously described with minor modifications[4]. Briefly, for each location reaction, approximately $4\times10^6$ cells were harvested followed by 10 min cross-linking with formaldehyde at a final concentration of 1% and washed twice in PBS. Cells were then lysed in Lysis Buffer (50 mM HEPES [pH7.4], 140 mM NaCl, 1% Triton X-100, 0.1% NaDeoxycholate, 1 mM EDTA supplemented with protease inhibitor cocktail (Roche, 11836170001)), sonicated to solubilise chromatin and shear the cross-linked DNA. Sonication was performed at 4°C with Biotuptor pico at default power for 40×30 second pulse (30 second pause between pulses). To retrieve chromatin-associated proteins, the whole cell extracts were incubated on a rotator overnight at 4°C with 10 μl magnetic Protein G Dynabeads (Invitrogen, 10004D) pre-coated with 2 μg of the indicated antibodies or Flag M2 beads (sigma, M8823). The beads were washed five times with Lysis Buffer, Lysis High Salt Buffer

(50 mM HEPES [pH7.4], 500 mM NaCl, 1% Triton X-100, 0.1% NaDeoxycholate, 1 mM EDTA), Wash Buffer (10 mM Tris [pH8.0], 250 mM LiCl, 0.5% NP40, 0.5% NaDeoxycholate, 1 mM EDTA) and TE buffer [pH8.0] two times each. Bound DNA-Immune-complexes were eluted off the beads in elution buffer (50 mM Tris [pH8.0], 1% SDS, 10 mM EDTA) by heating at 65$^{o}$C for 2 hours. DNA was purified using HighPure PCR Template Preparation Kit (Roche, 11796828001). Whole cell extracts were treated in parallel for cross-link reversal and DNA extraction. 200ng recovered DNA was used for each PCR reaction using Platinum$^{R}$*Taq* DNA Polymerase (Invitrogen, 10966). All reactions were performed in triplicate. Primer sets used to measure protein enrichment: CRISPR system (input, pair 22/23; 0.5 Kb, pair 6/7; 3.5 Kb, pair 8/9; 10 Kb, pair 18/19); DIvA system (0.5 Kb, pair 26/27; 1 Kb, pair 28/29; 2.5 Kb, pair 30/31; 5 Kb, pair 32/33

**Preparation of metaphase chromosomal spread**

Cells were plated in 60-mm dish and arrested in mitosis by 2-hour treatment with colcemid (final concentration of 200 ng/ml). Pre-warmed 75 mM KCl were then added to trypsinized cells and incubated for 15 min at 37°C. Then four drops of freshly prepared fixative (3:1 solution of methanol : acetic acid) were added. Cells were pelleted and resuspended in 5 ml fixative and incubated for 20 min at 4°C. After repeating the fixing step three or four times, cell pellets were resuspended in 0.5 ml fixative solution. Two or three drops of fluid were precipitated onto a pre-chilled slide from a height of 18 inches. Slides were air-dried thoroughly and stained using the Giemsa protocol. The mitotic chromosomes were observed and evaluated using an Olympus fluorescence microscope (BX51) at 1000× magnification.

**Cell survival assay**

For colony formation assays cells were plated at 500 cells each in 10 cm dish and cultured at 37$^{o}$C for 10-14 days, then fixed with methanol and stained with Giemsa solution. PARPi (Olaparib/KU0059436, Talazoparib/S7048, Selleck; Niraparib/MB5556, Veliparib/MB5524, Rucaparib/MB1643, Meilunbio) or Nutlin (ApexBio, A4228) was applied in the indicated concentration alone or simultaneously used with diluted Cisplatin (131102, Supertrack Bio-pharmaceutical). Cell proliferation of *WDR70* knockdown 293T cells were measured by cell counting every other day after transfection.

### Animals

All animal experiments were carried out in animal facilities of West China Second University Hospital or at outsourcing service (IDMO, Beijing) and were approved by local Ethics Committees and performed strictly in accordance with their ethics guidelines and regulations (Reference number, Medical Research 2018 (015)).

All mice were housed in accordance with approved protocols. All efforts were made to minimize suffering.

C57BL/6 mice were purchased from the animal facility of Sichuan University. *Trp53* knockout strain with deleted Exons 2-6 (B6.129S2-Trp53tm1/Nju) was purchased from Model Animal Research Centre of Nanjing University. Homozygous *Trp53*$^{null}$ was obtained from interbred *Trp53*$^{-/+}$ and genotyped by PCR. To obtain si*Wdr70* mice, *in vivo* knockdown of *Wdr70* was performed by injecting specific double strand siRNA (2.5 mg/Kg, si*Wdr70-001* (siG1117130927, target sequence: 5'-CUGCCAGAAUGGAAGCAUA-3', Ribobio, Guangzhou, China) and 25 µl RNA transfection reagent (Biotool, B45215) via tail vein twice a week. Injection started from 6 week after birth on a weekly basis. $_{Albumin}$*HBx* transgenic mice were provided by Cyagen. 465-bp HBx ORF is placed downstream of liver-specific *Albumin* promoter in pRp.Des3d vector. 1-5 ng/µl linearized pRP[Exp]-ALB>HBx:IRES:EGFP was micro-injected into pronuclei obtained from superovulated C57BL/6 and cultured overnight till 2- cell stage, followed by implantation into pseudopregnant female mice to generate $_{Albumin}$*HBx* transgenic pups. Liver-specific expression and maintenance of *HBx* was further confirmed by tail genotyping and RT-PCR (liver) of full length *HBx*. All experiments were carried out in heterozygotic $_{Albumin}$*HBx* mice capable of stable germline transmission after F1 generation.

For survival analysis on si*Wdr70*, *Trp53*$^{null}$ and double mutant mice, equivalent volume of PBS plus 25 µl RNA transfection reagents (si*Scramble* or si*Wdr70*) were injected into C57BL/6 and *Trp53*$^{null}$ group. Life span of individuals was determined according to the sacrifice date of terminal illness.

**PARP inhibition using nude mice and xenograft model**

Athymic nude immunodeficient mice (BALB/c nu/nu, SPF) were purchased from the animal center of Sichuan University. For T43 xenograft mouse models, female nude mice starting at 4-5 weeks of age were used for the experiments. To assess the tumorigenicity of L02 or T43 cells *in vivo*, mice were subcutaneously inoculated in both sides of armpits or hind flanks ($4\times10^6$ cells *per* site). When T43 xenografts had reached an average volume of approximate 0.01 $cm^3$, animals were randomized (using random number table) into treatment groups and five animals were selected for each group. For mono-treatment of Olaparib, each animal was administered intraperitoneally at dosage of 131.5 mg $kg^{-1}$ of body weight per day. For double treatment, Olaparib was administered at dosage of 41 mg $kg^{-1}$ of body weight per day, and Cisplatin at 0.42 mg $kg^{-1}$ of body weight every two days, respectively. A parallel group of mice was administered with PBS-diluted DMSO as control. Inoculated and drug-administered mice were observed each day. Sizes of tumours was measured by Vernier calliper and tumour volumes calculated by the three-dimensional measurement (length×width×height×π/6) until termination of the experiments. For excluding Outliers of tumour volume, Grubbs' test was applied (Gi>GO. 95, two-tailed test). Experiments were performed blindly.

PDX assays was established and performed in SPF facilities of Beijing IDMO Co, Ltd. Primary HBVHCC tissues were enzymatically dissociated and primary tumor cells were subcutaneously inoculated into the flank of NOD-Prkdcscid-IL2rg(em1 IDOM) (NPI) recipient mice. After sufficient tumor growth, tumors were passaged or cryopreserved for banking. For PARPi treatment of HBVHCC, 4 Cisplatin-insensitive HBVHCC and 1 HBV-free HCC were inoculated subcutaneously to NSG mice at 4-6 weeks of age. Randomized and age-matched males were used with an even split between control and drug administration. In total, 22 tumor-burdened mice survived to terminal stage of the experiments and data points were collected from these animals. Each tumor included a vehicle and an experimental group, which were allocated 2 - 3 repeats and performed blindly. Vehicle (12.5% DMSO in PBS) or combined Olaparib (33.3 mg/kg/day) + Cisplatin (3 mg/kg/2 days) treatment was administered by intraperitoneal injection. The health of the animal was monitored daily throughout therapy. Sizes of xenograft implantations were regularly measured by Vernier calliper and calculated for 3-dimentional volumes. Mean percentages of inhibition were calculated by %Inhibition = (mean(C)-mean(T))/mean(C) * 100%. HBV-postive and HBV-negative HCC materials used in PDX experiments were selected according to the HBV infection status: HBV+ tumors were serologically HBsAb- HBsAg+ HBcAb+ HBeAb+ HBeAg-, and HBV- tumors were HBsAb+ HBsAg- HBcAb- HBeAb- HBeAg-. All HCC were from Chinese male patients.

**Data analysis**

The percentage of TGI is defined as (1 - (mean volume of treated tumours)/(mean volume of control tumours)) x 100%.PFS (progression-free survival) was defined in this study as the condition where tumour volumes were under 300 $mm^3$. Mice with tumour sizes exceeding 300 $mm^3$ were assigned “progressive disease” and taken off PFS calculation. These mice could be enrolled again into PFS calculation if tumour regressed < 300 $mm^3$ during later treatment. The PFS curves were generated by the Kaplan–Meier method and log-rank tests were used to compare PFS in each experimental group.

**Statistics**

All histograms were presented as means±standard deviation. For quantitative analysis including ChIP assay, image analysis and repair analysis, at least three independent experiments were carried out. The Kaplan-Meier plot and the *log-rank* test were computed by SPSS 16.0 software to compare the survival outcomes between three groups. Other significant difference or *p* values between two groups was performed by

Two-tailed Student’s t tests using GraphPad Prism 6. End-point values of cell survival and anti-tumour assays were used for statistical analysis.

**Patient consent**

De-identified tumour material was collected from patients with solid tumours in agreement with local institutional ethical regulations and approved by local ethics board. Patient consent for tissue acquisition was obtained under the guidelines of each institution.

**Table S1. Peptide identification by mass spectrometry for spWdr70-interacting proteins**

| **Gene names** | **Coverage** (%) | **MW** (kDa) | **Description** |
|---|---|---|---|
| *wdr70/SPAC343.17c* | 90.55 | 63.6 | Uncharacterized WD repeat-containing protein |
| *rpn2/SPBC17D11.07c* | 57.68 | 107.2 | 19S proteasome regulatory subunit |
| *rpt6/SPBC23G7.12c* | 66.61 | 45.0 | 19S proteasome base subcomplex ATPase subunit |
| *rpt3/SPCC576.10c* | 39.93 | 43.5 | 19S proteasome base subcomplex ATPase subunit |
| *rpt4/SPCC1682.16* | 55.83 | 43.6 | 19S proteasome base subcomplex ATPase subunit |
| *rpn7/SPBC582.07c* | 57.03 | 46.7 | 19S proteasome regulatory subunit |
| *rpn5/SPAC1420.03* | 34.57 | 51.6 | 19S proteasome regulatory subunit |
| *rpn6/SPAC23G3.11* | 23.71 | 47.3 | 19S proteasome regulatory subunit |
| *rpt5/SPAC3A11.12c* | 37.38 | 48.8 | 19S proteasome base subcomplex ATPase subunit |
| *rpt1/SPBC16C6.07c* | 35.84 | 48.9 | 19S proteasome base subcomplex ATPase subunit |
| *rpt2/SPBC4.07c* | 10.04 | 50.0 | 19S proteasome base subcomplex ATPase subunit |
| *rpn3/SPBC119.01* | 47.06 | 57.3 | 19S proteasome regulatory subunit |
| *rpn1/SPBP19A11.03c* | 48.33 | 97.9 | 19S proteasome regulatory subunit |
| *pcu4/ SPAC3A11.08* | 1.5 | 85.3 | Cullin-4 |
| *rpn9/SPAC607.05* | 4.99 | 43.4 | 19S proteasome regulatory subunit |
| *rpn1302/SPCC16A11.16c* | 15.46 | 43.9 | 19S proteasome regulatory subunit Rpn13b |
| *hht1 / SPAC1834.04* | 5.15 | 15.3 | histone H3 h3.1 |
| *htb1 / SPCC622.09* | 9.52 | 13.8 | histone H2B |

### Table S2. Plasmids used in this study

| Plasmids | Source |
|---|---|
| pcDNA3-HA-HBx | Previous study |
| pCMV-Flag-WDR70 | Previous study |
| pLVX-puro-HA-HBx | Previous study |
| pLVX-shWDR70 | Purchased from Genechem |
| pLVX-sh53BP1 | Purchased from Genechem |
| pCMV-NHEJ | Gift from Prof. Jun Chen, Zhejiang University |
| pCMV-HR | Gift from Prof. Jun Chen, Zhejiang University |
| pCMV-SSA | Gift from Prof. Jun Chen, Zhejiang University |
| pMC1-1-p84-g1 | gRNA vector for *PPP1R12C/p84* locus, Purchased from Viewsolid biotech |
| pMC1-1-p84-g0 | gRNA vector (empty), Purchased from Viewsolid biotech |
| pLVX-G-BRCA1 | This study |
| pLVX-Flag-PSMD1 | This study |
| pLVX-Flag-PSMD2 | This study |
| pLVX-Flag-PSMD3 | This study |
| pLVX-Flag-PSMD4 | This study |
| pLVX-Flag-PSMD5 | This study |
| pLVX-Flag-PSMD6 | This study |
| pLVX-Flag-PSMD7 | This study |
| pLVX-Flag-PSMD8 | This study |
| pLVX-Flag-PSMD9 | This study |
| pLVX-Flag-PSMD10 | This study |
| pLVX-Flag-PSMD11 | This study |
| pLVX-Flag-PSMD12 | This study |
| pLVX-Flag-PSMD13 | This study |
| pLVX-Flag-POH1 | This study |
| pLVX-Flag-PSMC1 | This study |
| pLVX-Flag-PSMC2 | This study |
| pLVX-Flag-PSMC3 | This study |
| pLVX-Flag-PSMC4 | This study |
| pLVX-Flag-PSMC5 | This study |
| pLVX-Flag-PSMC6 | This study |
| pLVX-Flag-PSMD4-dUIM | This study |
| pFastBacHTA-6His-WDR70(112-654 aa) | This study |
| pLVX-Flag-ADRM1 | This study |
| pLVX-Flag-ADRM1-I75RF76RD79N | This study |
| pLVX-Flag-ADRM1-Pru-d | This study |
| pLVX-Flag-ADRM1-K99R | This study |
| pLVX-Flag-ADRM1-K99-only | This study |
| pET28a-ADRM1-Flag | This study |
| pET28a-ADRM1-K99RFlag | This study |
| pFastBacHTA-6His-DDB1 | This study |
| pFastBacHTA-6His-ROC1(5-108 aa) | This study |
| pFastBacHTA-6His-CUL4A(38-759 aa) | This study |
| pcDNA3-4HA-ubiquitin | This study |
| pcDNA3-4HA-ubiquitin-K48R | This study |
| pcDNA3-4HA-ubiquitin-K63R | This study |

**Table S3. Primers used in this study**

| | Primers | Application |
|---|---|---|
| Primer 1 | ATCATGGCCGACAAGCAGAAGAACG | efficiency of DSB repair, Normalize, forward |
| Primer 2 | CGGCGGCGGTCACGAACTCC | efficiency of DSB repair, Normalize, reverse |
| Primer 3 | TGACCACCCTGACCTACG | efficiency of DSB repair, HR and SSA repair, forward |
| Primer 4 | CACCTTGATGCCGTTCTTCTGC | efficiency of DSB repair, repair, reverse |
| Primer 5 | TCGGAGCAAGCTTGATTTAGGTGA | efficiency of DSB repair, NHEJ repair, forward |
| Primer 6 | CTAACTTTGGCTCTTCACCT | Amplicon at 0.5 Kb, forward |
| Primer 7 | GATGGAGAAAGAGAAAGGGA | Amplicon at 0.5 Kb, reverse |
| Primer 8 | TCGCCAGTGCTTTTTCTTTT | Amplicon at 3.5 Kb, forward |
| Primer 9 | GTTGGGGGATGATGAAAATG | Amplicon at 3.5 Kb, reverse |
| Primer 10 | GGCTGGAGTGCAATGGCATG | Amplicon at 5 Kb, forward |
| Primer 11 | AGACCAGCCTGGTCAACAT | Amplicon at 5 Kb, reverse |
| Primer 12 | TCCTGCAGAAATTGCTCATAAC | Amplicon at 6 Kb, forward |
| Primer 13 | ACGGCTGAGGGTCTTTCCAGT | Amplicon at 6 Kb, reverse |
| Primer 14 | GCCAACCTGACCAACATGG | Amplicon at 7 Kb, forward |
| Primer 15 | AGTGGCACGATCTTGGCTCA | Amplicon at 7 Kb, reverse |
| Primer 16 | GACCAGCCTGGCCAACATG | Amplicon at 8 Kb, forward |
| Primer 17 | CTGTTGCCCAGGCTGGAGTG | Amplicon at 8 Kb, forward |
| Primer 18 | ATGGCTCATGCCTGTAATCC | Amplicon at 10 Kb/ Total DNA, forward |
| Primer 19 | CAGCCTCCCAAGTAGCTGAG | Amplicon at 10 Kb/ Total DNA, reverse |
| Primer 20 | CCTCCCAGAGAACAAACAGC | Amplicon at 50 Kb/ Input, forward |
| Primer 21 | GGTTCGGGTAGGTTTTTCCT | Amplicon at 50 Kb/ Input, reverse |

| | | |
|---|---|---|
| Primer 22 | GCTCAGCTAGTCTTCTTCCTC | Amplicon for Uncut, forward |
| Primer 23 | CTTAGAGGTTCTGGCAAGGAG | Amplicon for Uncut, reverse |
| Primer 24 | GACGACGATAAGGAATTCATGGATTTATCTGCTCTTCG CGTTG | BRCA1 to pLVX-G, forward |
| Primer 25 | TAGTCTCGAGGAATTCTCAGTAGTGGCTGTGGGGGA TCTGGGG | BRCA1 to pLVX-G, reverse |
| Primer 26 | CCTGGATATGAGTTTGATCAGC | Amplicon at 0.5 Kb, DIvA system, forward |
| Primer 27 | CTCTCCTTTCGCTGACACTG | Amplicon at 0.5 Kb, DIvA system, reverse |
| Primer 28 | AGGAATTGACTGCGGTGTTC | Amplicon at 1 Kb, DIvA system, forward |
| Primer 29 | GGGGAGGAGGAAAGGTGTAG | Amplicon at 1 Kb, DIvA system, reverse |
| Primer 30 | GCCATAACAGAGGGTGGAAA | Amplicon at 2.5 Kb, DIvA system, forward |
| Primer 31 | AACTTTAGGATGGGGCTGCT | Amplicon at 2.5 Kb, DIvA system, reverse |
| Primer 32 | CAACATCCCTGATGACTACAGAC | Amplicon at 5 Kb, DIvA system, forward |
| Primer 33 | GGCAATAATGTTGCCTGCAA | Amplicon at 5 Kb, DIvA system, reverse |
| Primer 34 | GGCATCAGCAAAGTCTGAGC | Amplicon at α site of *p16* locus, forward |
| Primer 35 | CTGGGAGACAAGAGCGAAAC | Amplicon at α site of *p16* locus, reverse |
| Primer 36 | AGGGGAAGGAGAGAGCAGTC | Amplicon at β site of *p16* locus, forward |
| Primer 37 | GGGTGTTTGGTGTCATAGGG | Amplicon at β site of *p16* locus, reverse |
| Primer 38 | GGTAGGGAGTTCGAGACCAG | Amplicon at *GAPDH* locus, forward |
| Primer 39 | TCAACGCAGTTCAGTTAGGC | Amplicon at *GAPDH* locus, reverse |
| Primer 40 | CACCGAATAGTTACGGTC | Expression for *p16*, forward |
| Primer 41 | GCACGGGTCGGGTGAGAGTG | Expression for *p16*, reverse |
| Primer 42 | GACAAACGACCCCGTGAACTAC | Expression for *PSMD1*, forward |
| Primer 43 | CAAGGACGCCAACCACAGAAG | Expression for *PSMD1*, reverse |
| Primer 44 | CTCATCTCTGTTTCAAATCCAC | Expression for *PSMD2*, forward |

| | | |
|---|---|---|
| Primer 45 | GGCATGATATTGAGCTAACTG | Expression for *PSMD2*, reverse |
| Primer 46 | GGGAGAAGTTTCAAGCAGATG | Expression for *PSMD3*, forward |
| Primer 47 | TCTCGGGTGGAATAGATGTC | Expression for *PSMD3*, reverse |
| Primer 48 | GGATTGCTACGACTGGGACTG | Expression for *PSMD4*, forward |
| Primer 49 | CCTGCATCACGTCGTAATCATC | Expression for *PSMD4*, reverse |
| Primer 50 | GCTTGCTTATGAGAATAGGAC | Expression for *PSMD5*, forward |
| Primer 51 | GGCAGCACAGTGTAGTTCAGG | Expression for *PSMD5*, reverse |
| Primer 52 | CATGCATACAGTCAGCTGCTGG | Expression for *PSMD6*, forward |
| Primer 53 | GTTCTTGCTATCAGGTCTGTTG | Expression for *PSMD6*, reverse |
| Primer 54 | CCACCAGATCATCTACCAGCTGCA | Expression for *PSMD7*, forward |
| Primer 55 | GCATCCCGGTTGGCAATCTTG | Expression for *PSMD7*, reverse |
| Primer 56 | CCTTCTTCATTGACATCCTGCTCG | Expression for *PSMD8*, forward |
| Primer 57 | GTTCTGTGGAGGGAATGGTGGTG | Expression for *PSMD8*, reverse |
| Primer 58 | TCTGCAAGTGGATGATGAG | Expression for *PSMD9*, forward |
| Primer 59 | CGTGTTGGAACAAGTCTAAG | Expression for *PSMD9*, reverse |
| Primer 60 | GCTAATCCAGATGCTAAGGACC | Expression for *PSMD10*, forward |
| Primer 61 | GTAAATACTTGCTCCTTGGGAC | Expression for *PSMD10*, reverse |
| Primer 62 | CCAGAGTACAGATTGAACAC | Expression for *PSMD11*, forward |
| Primer 63 | GTTTCCAGAGCAGCTTCGTAAG | Expression for *PSMD11*, reverse |
| Primer 64 | CTTGAGAGTCCTGCAACGGATG | Expression for *PSMD12*, forward |
| Primer 65 | CCTGCTAATCTGTCTACTTTAGC | Expression for *PSMD12*, reverse |
| Primer 66 | GTTGTGCCTCATGGAGATGACT | Expression for *PSMD13*, forward |
| Primer 67 | GTCATGTGGACTCGTTTGTC | Expression for *PSMD13*, reverse |
| Primer 68 | GACACTTCAGGACTACAGTGAAC | Expression for *PSMD14*, forward |

| | | |
|---|---|---|
| Primer 69 | GAGGTCATAAGTACATCCACATG | Expression for *PSMD14*, reverse |
| Primer 70 | CTTTGGATCCAGCACTTATCAG | Expression for *C1*, forward |
| Primer 71 | CATCAGACCAGCTTCTGTACAG | Expression for *C1*, reverse |
| Primer 72 | CAGACCTGATACTTTGGATCC | Expression for *C2*, forward |
| Primer 73 | CTCTGATGGCAAACATACCAG | Expression for *C2*, reverse |
| Primer 74 | CTTTGACAGTGAGAAGGCTGG | Expression for *C3*, forward |
| Primer 75 | GTAGTTCACGTCAGGACTGAC | Expression for *C3*, reverse |
| Primer 76 | CTTCATAGACGAGATTGATGCC | Expression for *C4*, forward |
| Primer 77 | CATCTTGCTAGTGATAGTGGAG | Expression for *C4*, reverse |
| Primer 78 | GATTGATATCCTGGACTCGGCAC | Expression for *C5*, forward |
| Primer 79 | CTCAAAGTCCTCCTGAGTGACATG | Expression for *C5*, reverse |
| Primer 80 | GATCATGGCTACAAACAGACCAG | Expression for *C6*, forward |
| Primer 81 | CATGATCAGCACGAATTGCGAAC | Expression for *C6*, reverse |
| Primer 82 | ACACTGTTACAAGAGTCCGACCACTG | Amplicon at 1 Kb, resection assay, forward |
| Primer 83 | GCATATGCTGCCCAACTTTGGCTAATTC | Amplicon at 1 Kb, resection assay, reverse |
| Primer 84 | ACCCTACAGTAGTACTAAGAAGGAC | Amplicon at 3.3 Kb, resection assay, forward |
| Primer 85 | ATCTGGCTAGGAAACATTAGTTTAATATTC | Amplicon at 3.3 Kb, resection assay, reverse |
| Primer 86 | GATTGGCTGTGGAGTGTGACTCA | Amplicon for total DNA, resection assay, forward |
| Primer 87 | ATCCTCCATCTTGTCCCCTT | Amplicon for total DNA, resection assay, reverse |

**Table S4. Antibodies used in this study**

| Antibodies | Source | dilutions |
|---|---|---|
| Rabbit anti-phosphor-Serine 33, RPA32 | NOVUS, NB100-544 | 1:3000 |
| Mouse anti-α-Tubulin | Sigma, T6074 | 1:5000 |
| Rabbit anti-RAD51 | Proteintech, 14961-1-AP | 1:500 |
| Rabbit anti-53BP1 | Bethyl, A300-272A | 1:1000 |
| Rabbit anti-phosphor-Serine 1524, BRCA1 | Bethyl, A300-001A | 1:500 |
| HRP-conjugated anti-mouse IgG | DAKO, P0260 | 1:2000 |
| HRP-conjugated anti-rabbit IgG | DAKO, P0448 | 1:3000 |
| FITC- conjugated anti-mouse IgG | Sigma, F0257 | 1:300 |
| CY3- conjugated anti- rabbit IgG | Sigma, C2306 | 1:300 |
| Rabbit anti-BRCA1 | Millipore, 07-434 | 1:500 |
| Rabbit anti-Ki67 | Abcam, ab16667 | 1:200 |
| Mouse anti-γH2AX | Millipore, 05-636 | 1:500 |
| Rabbit anti-Flag | HuaBio, custom made | 1:2000 |
| Mouse anti-PSMD5 | Abnova，H00005711-M01 | 1:500 |
| Rabbit anti-MRE11 | CST, ,4895S | 1:500 |
| Rabbit anti-WDR70 | Bethyl, A301-871A | 1:500 |
| Rabbit anti-DDB1 | Epitomics, 3821-1-1 | 1:500 |
| Rabbit anti-Strep | HuaBio, HA500061 | 1:2000 |
| Rat anti-HA | Roche, 11867423001 | 1:500 |
| Rabbit anti-GST | ZENBIO, 300195 | 1:2000 |
| Rabbit anti-ADRM1 | Proteintech, 11468-1-AP | 1:500 |
| Mouse anti-H3 | Millipore, 05-1341 | 1:20000 |
| HRP-conjugated streptavidin | Sigma, S2438 | 1:3000 |

**Table S5. siRNA used in this study**

| Gene | Sequence | Source |
|---|---|---|
| *siWDR70* | 5’-CUGCCAGAAUGGAAGCAUA-3’ | Ribobio |
| *siDDB1* | 5’-CCUGUUGAUUGCCAAAAAC-3’ | Ribobio |
| *si53BP1* | 5’-GGCCUUUGCCUCUCAACAA-3’ | Ribobio |
| *siBRCA1* | 5’-GGAACCUGUCUCCACAAAG-3’ | Ribobio |
| si*PSMD1* | 5’-GAGGCAUCAUCAUUCUGAA-3’ | Ribobio |
| si*PSMD2* | 5’-GAAUGCUGGUUACGUUUGA-3’ | Ribobio |
| si*PSMD3* | 5’-GCCGCAAAGUGUUACUAUU-3’ | Ribobio |
| si*PSMD4-1* | 5’-GCACCGACAAGGCAAGAAU-3’ | Ribobio |
| si*PSMD4-2* | 5’-GGCGGAAUCAGCAGACAUU-3’ | Ribobio |
| si*PSMD5-1* | 5’-GGAUGACAGAAUCCUGGUU-3’ | Ribobio |
| si*PSMD5-2* | 5’-GCGACAAUAUCUUGCUCAA-3’ | Ribobio |
| si*PSMD6* | 5’-GAGCGAAAUUCGCGAUGCA-3’ | Ribobio |
| si*PSMD7* | 5’-UGGUCAUCAUUGAUGUGAA-3’ | Ribobio |
| si*PSMD8* | 5’-GCAUGUACGAGCAACUCAA-3’ | Ribobio |
| si*PSMD9* | 5’-CCAGCUUAGACUUGUUCCA-3’ | Ribobio |
| si*PSMD10* | 5’-AGAGUAUUCUGGCCGAUAA-3’ | Ribobio |
| si*PSMD11* | 5’-GCCUUAACUUCUGCUCGAA-3’ | Ribobio |
| si*PSMD12* | 5’-GUACUUAUGUUGAGGAAAU-3’ | Ribobio |
| si*PSMD13* | 5’-GGUUCACAGUCGUUUCUAU-3’ | Ribobio |
| si*POH1-1* | 5’-AUACCGUCAGAGUGAUUGA-3’ | Ribobio |
| si*POH1-2* | 5’-AAGGCCGGAGAUGGUUGUU-3’ | Ribobio |
| si*PSMC1* | 5’-GGAGACCUAUGCAGAUAUU-3’ | Ribobio |
| si*PSMC2* | 5’-GGUCAGAGCACUUACUCUA-3’ | Ribobio |
| si*PSMC3* | 5’-GGAGGAUGGUGCCAAUAUU-3’ | Ribobio |
| si*PSMC4* | 5’-GGAAGACCAUGUUGGCAAA-3’ | Ribobio |
| si*PSMC5* | 5’-GGAACGAACUAAAUGCUAA-3’ | Ribobio |
| si*PSMC6* | 5’-GUUCUAUUGUAGACAAGUA-3’ | Ribobio |
| siRNF8 | 001: 5’-GGACAAUUAUGGACAACAA-3’<br>002: 5’-GGACGAGGAUUUGGUGUCA-3’<br>003: 5’-GCUAGAGAAUGAGCUCCAA-3’ | Ribobio |
| siRNF168 | 001: 5’-GCAGUCAGUUAAUAGAAGA-3’<br>002: 5’-GUGGAACUGUGGACGAUAA-3’<br>003: 5’-CCUACAGCCUAGCAUUUCA-3’ | Ribobio |
| siMRE11 | 001: 5’-GCCTCGAGTTATTAAGAAA-3’<br>002: 5’-GGATATTGTTCTAGCTAAT-3’<br>003: 5’-GGAAATGATACGTTTGTAA-3’ | Ribobio |
| siRNF20 | 001: 5’- GCUAAACAGUGGAGAUAAU -3’<br>002: 5’- CCAAUGAAAUCAAGUCUAA -3’<br>003: 5’- GGAGAAGGAUGAUGCAAAU -3’ | Ribobio |
| GlaNAC-siENDOD1 | 5’-/i2FG*/i2OMeC*/i2FA/i2OMeA/i2FG/i2OMeC/i2FG/i2OMeG/i2FA/i2FU/<br>i2FU/i2OMeG/i2FG/i2OMeC/i2FU/i2OMeA/i2FC/i2OMeA/i2FA-3’-GalNac3 | Ribobio |
